# Endogenous responses in brain pH and *P*_O2_ in a rodent model of birth asphyxia

**DOI:** 10.1101/848291

**Authors:** Alexey S. Pospelov, Martin Puskarjov, Kai Kaila, Juha Voipio

**Author notes:** Correspondence: Juha Voipio, Faculty of Biological and Environmental Sciences, Molecular and Integrative Biosciences, PO Box 64, FI-00014 University of Finland, Finland.

## Abstract

**Aim:** To study brain-sparing physiological responses in a rodent model of birth asphyxia which reproduces the asphyxia-defining systemic hypoxia and hypercapnia.

**Methods:** Steady or intermittent asphyxia was induced for 15-45 min in anesthetized 6 and 11 days old rats and neonatal guinea pigs using gases containing 5% or 9% O_2_ plus 20% CO_2_ (in N_2_). Hypoxia and hypercapnia were induced with low O_2_ and high CO_2_, respectively. Oxygen partial pressure (*P*_O2_) and pH were measured with microsensors within the brain and subcutaneous (“body”) tissue. Blood lactate was measured after asphyxia.

**Results:** Brain and body *P*_O2_ fell to apparent zero with little recovery during 5% O_2_ asphyxia and 5% or 9% O_2_ hypoxia, and increased more than twofold during 20% CO_2_ hypercapnia. Unlike body *P*_O2_, brain *P*_O2_ recovered rapidly to control after a transient fall (rat), or was slightly higher than control (guinea pig) during 9% O_2_ asphyxia. Asphyxia (5% O_2_) induced a respiratory acidosis paralleled by a progressive metabolic (lact)acidosis that was much smaller within than outside the brain. Hypoxia (5% O_2_) produced brain-confined alkalosis. Hypercapnia outlasting asphyxia suppressed pH recovery and prolonged the post-asphyxia *P*_O2_ overshoot. All pH changes were accompanied by consistent shifts in the blood-brain barrier potential.

**Conclusion:** Regardless of brain maturation stage, hypercapnia can restore brain *P*_O2_ and protect the brain against metabolic acidosis despite compromised oxygen availability during asphyxia. This effect extends to recovery phase if normocapnia is restored slowly, and it is absent during hypoxia, demonstrating that exposure to hypoxia does not mimic asphyxia.

## 1 INTRODUCTION

Severe birth asphyxia (BA; also known as perinatal asphyxia) is the main cause of disability and mortality of human neonates worldwide, with more than one million casualties annually.^1^ The number of the surviving, afflicted individuals is not known, but there is reason to believe that it is much higher. Thus, BA makes a significant contribution to the total burden of disease in human populations, based on aberrant development and dysfunctions of organs, which are highly reliant on oxidative energy metabolism, especially the brain. The immediate pathological effect of BA on the brain manifests as hypoxic-ischemic encephalopathy (HIE), and the lifelong outcomes of HIE include a wide spectrum of psychiatric and neurological diseases and disorders, including cognitive defects, autism, epilepsy and cerebral palsy.^2–6^

Therapeutic hypothermia is currently the only generally accepted treatment for near-term and term newborns with moderate or severe BA, but it provides incomplete neuroprotection.^7–11^ In addition to HIE, BA causes a wide spectrum of (often causally connected) dysfunctions including those of the adrenals and the heart. Obviously, advances in understanding the basic physiology and pathophysiology of BA and the mechanisms that lead to HIE and adverse lifelong outcomes will promote development of more effective therapies.^12^

By definition, BA implies a decrease in systemic O_2_ (hypoxia) and an increase in CO_2_ (hypercapnia). Hypoxia and hypercapnia occur also during normal uncomplicated deliveries, and they play an essential role in triggering endogenous mechanisms that operate to centralise blood flow to critically oxygen-dependent organs.^13–15^ Augmenting endogenous neuroprotective mechanisms or supplementing their effectors has been frequently suggested as basis for novel therapeutic interventions for BA.^16, 17^

Regarding the translational value of animal models, understanding of the systems-physiological mechanisms involved in BA and related conditions^18^ has been significantly improved by elegant work on pathophysiological and intrinsic protective mechanisms in large-animal models such as sheep and pigs.^14–16^ However, these mechanisms have remained largely unexplored in standard laboratory rodents. Extending the systems-level work on BA from large animal models to laboratory rodents will offer the potential of studying the mechanistic aspects of BA and its consequences using the expanding array of neurobiological research methods available today, from molecules to systems, which have been largely tailored for rats and mice.

Our present experiments are mainly based on postnatal day (P) 6 and 11 rat pups which, in terms of neurodevelopmental (especially cortical) milestones, roughly correspond to preterm and full-term human neonates, respectively.^19–22^ Asphyxia is induced by applying an ambient gas mixture containing 5% O_2_ and 20% CO_2_ (balanced with N_2_) using two paradigms: *monophasic asphyxia* (“steady asphyxia”) which corresponds to an acute complication such as placental abruption or maintained cord compression, and *intermittent asphyxia* where the hypoxia is applied in repetitive steps at 5% O_2_ and 9% O_2_ (at constant 20% CO_2_) in order to roughly mimic the effects of recurring contractions during prolonged parturition. We use here infant rats of two ages, because even if a given insult would have similar immediate effects, the long-term outcomes in e.g. brain functions and behaviour (not studied presently) will depend on the stage of development of the brain at the time of insult.^19, 23^ We also included experiments on P0-2 guinea pigs to provide a more comprehensive translational basis and also because there is a recent increase in experimental work on early-life disorders in this species.^24, 25^

The main aim of our present work is to examine the complex and interconnected effects of BA on brain pH and oxygen partial pressure (*P*_O2_) levels. These are the two fundamental physiological variables which are strongly affected at the onset, during and after asphyxia. Notably, systemic acidosis (typically blood pH <7.0) is used as one of the standard diagnostic criteria of BA. Both pH and *P*_O2_ are known to be powerful modulators of the function of practically all organs and organ systems, including cardiorespiratory regulation^13, 26^ and the excitability of the immature brain.^27, 28^ Regarding the latter, a large variety of key molecules involved in neuronal signalling show a functionally synergistic (most likely evolutionary-contingent)^29^ sensitivity to pH, whereby brain acidosis acts to suppress neuronal excitability while alkalosis has the opposite effect.^29–31^ This notion gets further impact in the context of BA from observations that experimental hypoxia as such (i.e., not as a component of asphyxia) leads to a gross increase in neuronal network excitability as has been shown in both *in vivo* and *in vitro* conditions.^32–36^ “Pure” hypoxia, i.e. hypoxia without simultaneous hypercapnia – which never takes place in natural conditions – has dominated thinking not only in translational research of hypoxic-ischemic brain damage (see Discussion) but also in clinical practice, particularly regarding methods of resuscitation.^37^ The differences between pure hypoxia and asphyxia are further underscored by the profound influence of CO_2_ on cerebral blood flow (CBF).^13^ Thus, understanding the physiological and pathophysiological consequences of BA, including the propensity for subsequent manifestation of HIE, requires information on i) the magnitude and time course of perturbations of acid-base regulation and oxygenation in brain tissue, and on ii) the adaptive mechanisms which act in a “brain-sparing” protective manner. Using pH- and O_2_-selective microsensors implanted into the brain and subcutaneous tissue (“body”), we show here that experimental BA produces a fast and large CO_2_-mediated (respiratory) acidosis, which is combined to a slow metabolic acidosis that is much smaller within than outside brain tissue. A striking action by brain-sparing mechanisms was observed in simultaneous measurements of brain parenchyma and body *P*_O2_, which demonstrated a full restoration of the brain *P*_O2_ in response to steady and intermittent exposure to 9% O_2_ in the presence of 20% CO_2_.

This adaptive mechanism was absent under conditions of pure 9% hypoxia. Moreover, combined to the unexpected brain *alkalosis* which we observed immediately in response to pure hypoxia, these data show that exposure of an infant rodent to hypoxia^32, 38^ does not reproduce the physiological responses to asphyxia. Finally, we demonstrate that Graded Restoration of Normocapnia (GRN) following asphyxia, a putative therapeutic strategy,^39^ slows down the pro-excitatory recovery of brain pH and extends the duration of the post-asphyxia *P*_O2_ overshoot. Thus, in the neonatal intensive care unit, judicious tailoring of the GRN protocol would provide a possibility to block the frequent and deleterious post-BA hypocapnia.^40, 41^ Using GRN, it should also be possible to set the duration of the post-asphyxia *P*_O2_ overshoot to enhance brain oxygenation immediately after birth.^42, 43^

## 2 RESULTS

### 2.1 Changes in brain and body pH induced by asphyxia, hypercarbia and hypoxia in P6 rats

Birth asphyxia is associated with systemic acidosis that has a respiratory and a metabolic component due to CO_2_ accumulation and to O_2_ deficit, respectively. We first carried out a series of experiments on P6 rats in order to analyse changes in extracellular brain pH and in subcutaneous tissue pH (brain pH and body pH, respectively; see Materials and Methods). Recording of body pH using pH sensitive microelectrodes is continuous and therefore avoids problems associated with repeated blood sampling and it provides a continuous measure of systemic pH with good correlation with blood pH.^44^ Below, we also addressed separately the effects of hypoxia and hypercapnia, the two components of asphyxia, on brain and body pH.

#### Steady asphyxia

The mean brain pH at baseline in all experiments at P6 was 7.31 ± 0.01 (mean ± SEM, n = 32). Asphyxia (5% O_2_ and 20 CO_2_; for 45 min) caused immediately a rapid fall in brain pH, followed by a very slow and progressive acidification with a smaller amplitude (Figure 1A). The fall in pH within the first 10 min was 0.47 ± 0.01 (n = 6) with a maximum fall rate of 0.21 ± pH units per min, whereas the subsequent slow acidification was 0.10 ± 0.02 with a rate of 0.003 ± 0.0005 pH units per min, reaching a final pH of 6.75 ± 0.02 by the end of the 45 min asphyxia period.

**FIGURE 1.**
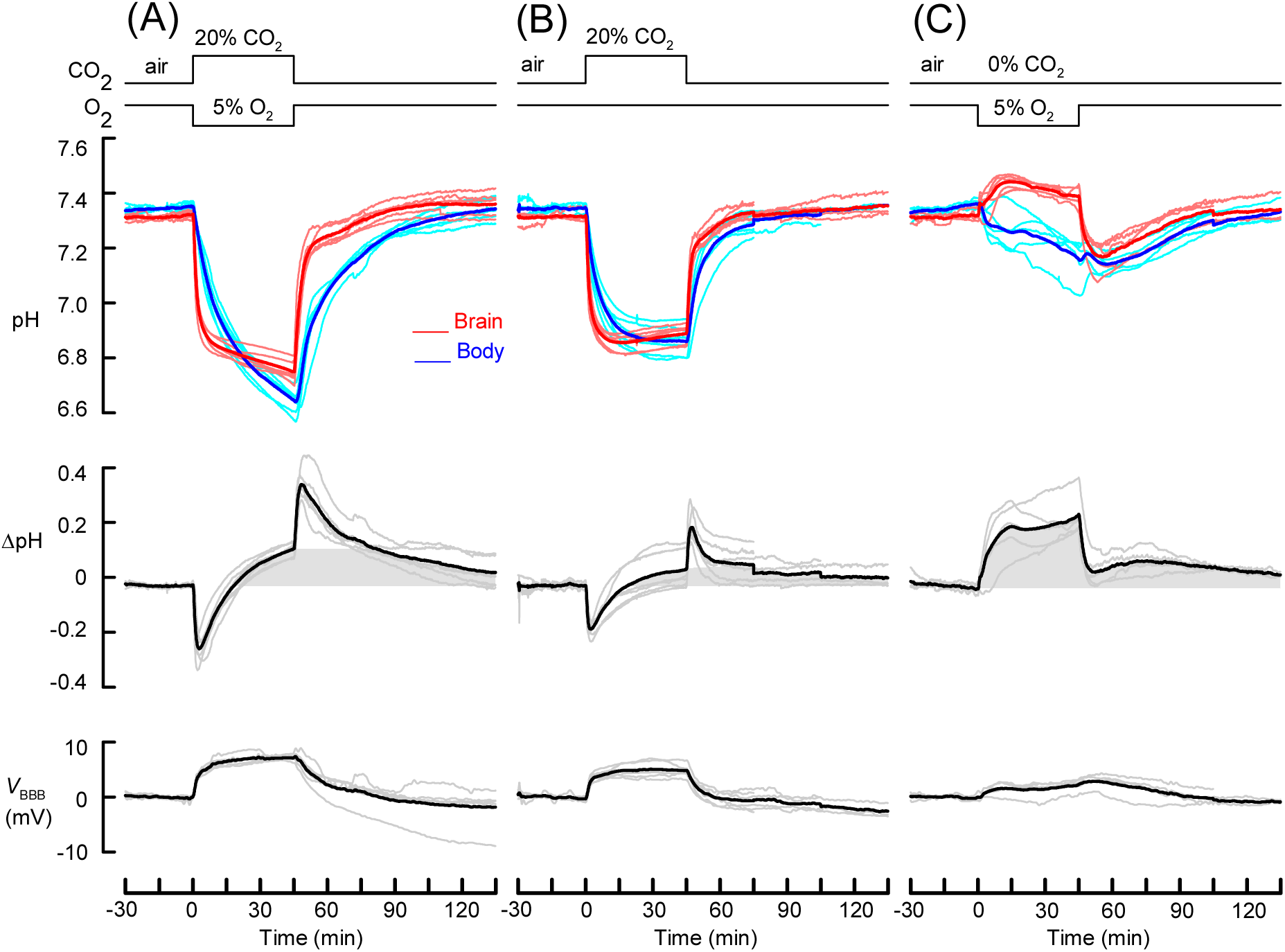
Experimental asphyxia, hypercapnia and hypoxia induced pH responses in P6 rats. **(A)** Brain (red) and body (blue) pH responses to 45 min 5% O_2_ asphyxia (top panel), their difference (middle panel) and the blood-brain barrier potential *V*_BBB_ (bottom panel). In this and subsequent Figures, superimposed light-coloured thin traces and dark-coloured thick traces show individual recordings and their mean, respectively. The line graphs above data traces indicate timing with which the animals were exposed to different gases. Grey shading is used to highlight the difference of brain and body pH, excluding the transient shifts caused by the slower respiratory body pH response upon changes in inhaled CO_2_. **(B)** Responses to 45 min hypercapnia induced by 20% CO_2_. **(C)** Responses to 45 hypoxia induced by reducing the O_2_ content in inhaled gas to 5%. In this and subsequent Figures some traces of individual recordings do not continue until the end of the recovery phase because recording was discontinued, which accounts for the small stepwise shifts seen in the mean traces.

The post-asphyxia recovery of brain acidosis was very fast (Figure 1A). It consisted of two phases, where the first 15 min was a mirror image of the asphyxia-induced rapid fall in pH, and it was followed by a second phase after about 20 min. Compared to brain pH changes seen previously upon a more moderate asphyxia (9% O_2_ and 20% CO_2_),^39^ the present results were different in that i) brain pH did not start to recover during asphyxia but decreased, albeit slowly, until the end of the asphyxia period, and ii) the final pH level at the end of the recovery period was only slightly higher than the initial baseline pH (by 0.05 ± 0.02, *P* = 0.034; paired t-test).

The mean body pH at baseline in all experiments at P6 was 7.34 ± 0.01 (n = 32). In contrast to the brain pH response during asphyxia, body pH changes did not have two kinetically distinct phases. Although initially slower than in the brain (maximum rates of acidosis in body and brain mean pH were 0.051 and 0.21 pH units per minute, respectively), the fall in body pH reached a higher amplitude of 0.70 ± 0.02 pH units at 45 min (n = 6; *P* = 0.00007, paired t-test; pH 6.65 ± 0.02). Also, the recovery of body pH was slower. The temporal behaviour of the difference (ΔpH = brain pH – body pH) is illustrated in the middle panel in Figure 1A. Notably, a slow shift in the alkaline direction of brain vs. body pH develops during asphyxia, which does not fully fade out by the end of the 90 min recovery.

#### Hypercapnia

Hypercapnia induced by 20% CO_2_ caused a prompt fall in brain pH with a maximum rate (0.21 ± 0.03 pH units per min) and amplitude (0.45 ± 0.01 at 10 min; n = 6; *P* = 0.294; Figure 1B) similar to what was seen in asphyxia. However, in contrast to asphyxia, there was no further slow acidosis but, instead, a small but consistent rise of 0.03 pH units after the peak acidosis of 6.85 ± 0.01 at 17.6 ± 1.5 min of hypercapnia (final pH at 45 min 6.89 ± 0.02). In contrast to asphyxia, hypercapnia led to a fall in body pH which attained a steady maximum amplitude that was almost identical to that generated simultaneously in the brain (0.48 ± 0.02, n = 6 vs. 0.43 ± 0.02 at 45 min, respectively; *P* = 0.099, paired t-test; final body pH at 45 min 6.84 ± 0.04). Notably, such pH changes are very similar to the “passive” (i.e., purely physicochemical) acid shift of 0.6 pH units that would be generated by an increase in CO_2_ from 5% to 20% in a physiological CO_2_/bicarbonate solution with no other buffers.^45^ This suggests that the rapid fall in brain pH, which constitutes a major part of brain acidosis during experimental asphyxia, is primarily caused by the increase in CO_2_ in the brain extracellular space where the non-CO2 buffer power is low.^46^ The faster responses in brain vs. body pH (see the transient “on” and “off” responses in the ΔpH traces in Figure 1) are readily accounted for by more effective perfusion of brain tissue compared to the subcutaneous site of body pH measurement.

Thus, striking and physiologically relevant differences were observed when comparing the neonates’ responses in body pH to asphyxia and hypercapnia. In particular, the above data indicate that the experimental asphyxia triggered a metabolic acidosis that is much larger in the body than in the brain.

While consistent with metabolic acidosis during the experimental asphyxia, the above dynamic pH data provide indirect evidence for its generation. Therefore, we took blood samples from P6 rats and, indeed, found that the present asphyxia protocol elevates blood lactate in only 15 min from 0.88 ± 0.15 mmol L^-1^ (n = 5) in controls to 11.1 ± 0.80 mmol L^-1^ (n = 5), see also ref. [^47^]. This result is of further (translational) importance given that generation of lactic acid is largely responsible for the base deficit (negative base excess) which is one of the key diagnostic criteria of BA.

#### Hypoxia

As a whole, the above data suggest that an endogenous mechanism protects the brain against metabolic acidosis and base deficit during asphyxia. This idea gained further support from the simultaneously measured brain and body pH responses to hypoxia (5% O_2_; Figure 1C). Unexpectedly, and in sharp contrast to what was seen during asphyxia, brain pH measurements demonstrated an *alkaline* shift upon hypoxia. Brain pH reached a maximum increase of 0.13 at 16 ± 1.5 min (7.44 ± 0.011 vs. baseline, n = 5; *P* = 0.0003, paired t-test) and, while losing amplitude, it remained above control throughout the 45 min hypoxia period (in 5 of 5 recordings). Return to air caused a transient rebound acidosis followed by recovery of brain pH to the control level. The alkaline pH response suggests that hypoxia triggers net extrusion of acid equivalents across the blood-brain barrier (BBB) at a rate that exceeds net acid production within the brain parenchyma.

The effective compartmentalization at the level of the BBB was also seen in measurements of body pH during hypoxia, in which a gradual monophasic acidification (amplitude at 45 min 0.18 ± 0.04, n = 5) with no initial alkalosis was seen. This is reminiscent of the dynamics of the slow (apparently metabolic, see above) body acidosis during asphyxia. Together, these responses resulted in a robust positive shift in ΔpH lasting throughout the hypoxia (Figure 1C). A comparable prolonged positive shift in ΔpH was evoked also by asphyxia but not by hypercapnia (see the shaded areas in Figure 1 that indicate the slowly-generated positive (brain more alkaline) ΔpH and where the ΔpH transients caused by step changes in inhaled CO_2_ have been excluded).

As a final conclusion based on the data in Figure 1, the time courses and amplitudes of brain and body pH changes related to asphyxia (Figure 1A) seem to behave roughly like sums of the pH responses triggered by the two underlying components, hypercapnia (Figure 1B) and hypoxia (Figure 1C), recorded in isolation.

### 2.2 pH-induced changes in the trans-BBB potential in P6 rats

The BBB maintains a pH-sensitive potential difference between brain tissue and the rest of the body (*V*_BBB_).^48, 49^ We monitored the *V*_BBB_ signal by measuring changes in the voltage between the brain and body (see Materials and Methods). The acid shifts induced by asphyxia or hypercapnia were tightly paralleled, as expected, by positive shifts in *V*_BBB_ with maximum amplitudes of 7.1 ± 0.18 mV (n = 6) and 4.9 ± 0.49 mV (n = 6), respectively (*P* = 0.0015). The response in *V*_BBB_ upon hypoxia was positive which, together with its smaller amplitude and time course, suggests a dependence on body pH, not on brain pH. Previous work shows that respiration-induced mV-level slow EEG shifts generated by the human BBB can be readily measured using non-invasive DC-coupled EEG.^49^ Thus, measuring DC-EEG shifts may open up a new window for brain monitoring during recovery from BA (see Discussion).

### 2.3 Brain and body *P*_O2_ changes upon asphyxia, hypercapnia and hypoxia in P6 rats

The results above raise questions about oxygen availability and consumption in brain vs. body during the experimental manoeuvres. Because CO_2_ is a well-known vasodilator,^13^ we next carried out tissue *P*_O2_ recordings in P6 rats.

The mean baseline levels of brain and body *P*_O2_ based in all experiments on the P6 rats were 20.8 ± 0.9 mmHg, n = 54, and 26.0 ± 1.8 mmHg, n = 23, respectively. These levels are much lower than i) *P*_O2_ in inhaled air, which is about 160 mmHg, and ii) arterial blood *P*_O2_ of 90 to 100 mmHg, which corresponds to normal 96% to 98% oxygen saturation,^50^ consistent with tissue *P*_O2_ levels reflecting a balance between O_2_ delivery and consumption.

The 5% O_2_ / 20% CO_2_ asphyxia resulted in a very rapid fall in both brain and body *P*_O2_ to stable, apparent zero levels (see Materials and Methods) that were maintained for the whole 45 min asphyxia period (Figure 2A; n = 8 and 4 for brain and body, respectively). During recovery, inhaling air evoked a large transient rise in brain and body *P*_O2_. Mean brain *P*_O2_ peaked in 2.9 ± 0.8 min at 68 ± 3.3 mmHg, and in the body in 6.6. ± 0.8 min at 49.2 ± 5.9 mmHg. The peaks were followed by a rapid fall in *P*_O2_ to hypoxic levels below baseline after ∼16 min and ∼23 min post-asphyxia in the brain and body, respectively. Near-control *P*_O2_ levels were restored by the end of the >90 min recovery period. Thus, the more transient nature of the rise in brain *vs.* body *P*_O2_ during recovery from asphyxia is in line with the more rapid kinetics of the CO_2_-induced brain acidosis (Figures 2A vs. 1B).

**FIGURE 2.**
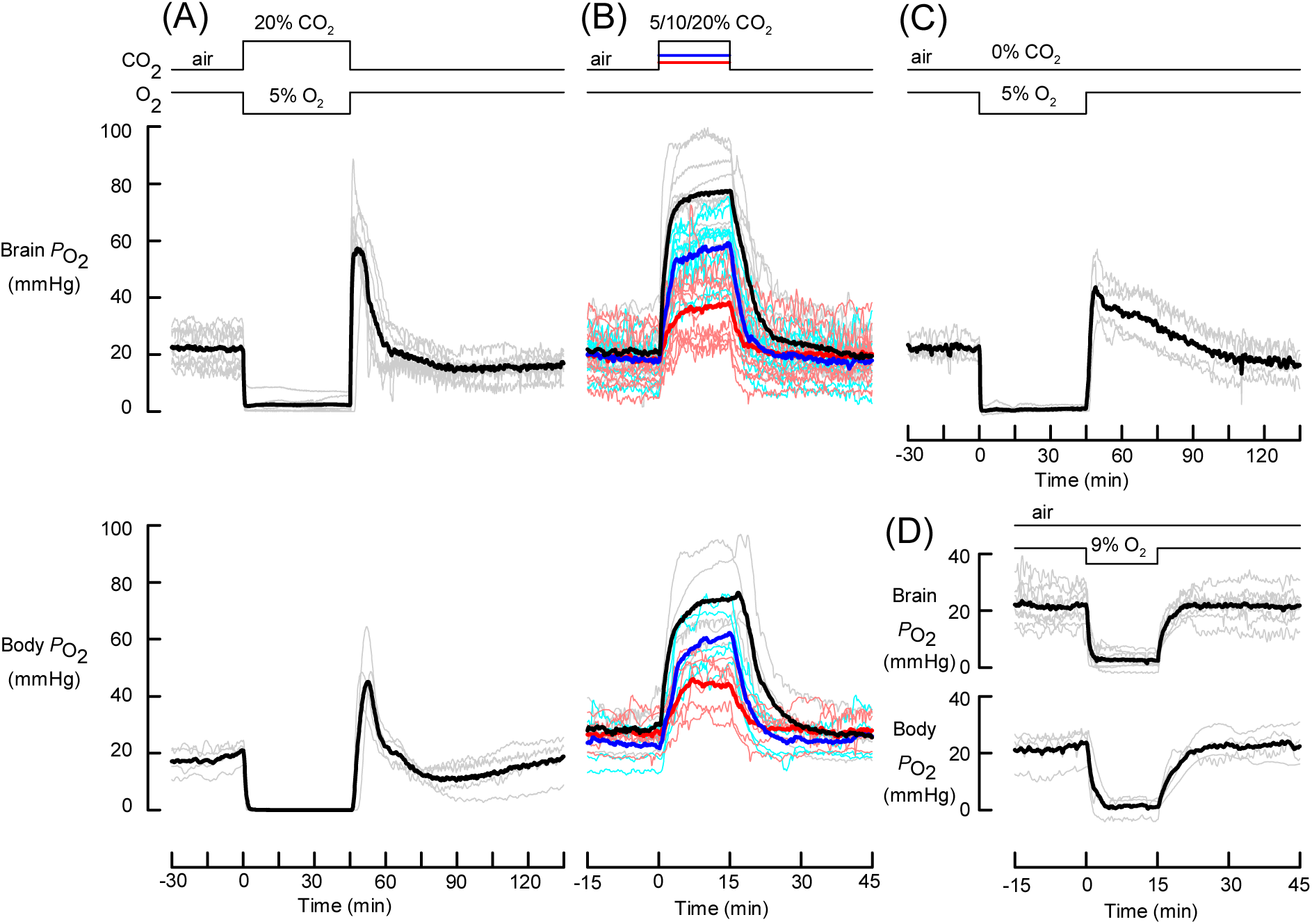
Experimental asphyxia, hypercapnia and hypoxia induced *P*_O2_ responses in P6 rats**. (A)** Brain (upper panel) and body (lower panel) *P*_O2_ fall rapidly to apparent zero during 45 min 5% O_2_ asphyxia and show a large transient overshoot during recovery in air. **(B)** Brain and body *P*_O2_ increase with increasing levels of hypercapnia that were generated by letting the rats inhale for 15 min 5%, 10% or 20% CO_2_ containing gas mixtures (red, blue and black traces, respectively) while keeping O_2_ at its normal level in air. At least 30 min was allowed for recovery between subsequent CO_2_ applications that were given in a pseudorandom order. **(C)** Brain *P*_O2_ falls rapidly to apparent zero when the animals are exposed to 5% O_2_ hypoxia for 45 min, and recovers with an overshoot of lower amplitude and longer duration compared to that seen in (A) after 5% O_2_ asphyxia. **(D)** A fall to apparent zero in both brain and body *P*_O2_ is seen even upon a more moderate hypoxia (9% O_2_ for 15 min).

The high *P*_O2_ values seen after asphyxia may reflect reduced O_2_ consumption and/or increased perfusion and O_2_ supply due to CO_2_-induced vasodilation outlasting the experimental asphyxia period. Therefore, we next exposed P6 rats to different levels of CO_2_ while maintaining ambient O_2_ at 20% throughout the experiments. Fifteen min periods of hypercapnia induced rapid and pronounced increases in brain and body *P*_O2_, both of which showed a monotonic dependence on inhaled CO_2_ (0 to 20% CO_2_; maximum increases in *P*_O2_ upon 15 min exposure to 5%, 10% and 20% CO_2_ were 19.7 ± 1.8 mmHg (n = 11), 38.9 ± 3.3 mmHg (n = 10) and 55.0 ± 3.2 mmHg (n = 10) in brain *P*_O2_, and 17.2 ± 2.3 mmHg (n = 6), 37.6 ± 3.5 mmHg (n = 5) and 45.2 ± 7.1 mmHg (n = 5) in body *P*_O2_; Figure 2B). These result show that the baseline brain and body *P*_O2_ levels are set by constraints in O_2_ supply and delivery (see above and Discussion). Notably, the level and time course of brain *P*_O2_ response after 20% CO_2_ exposure showed striking resemblance to what was seen after asphyxia. This suggests that the hypercapnia which is associated with asphyxia acts to reduce tissue hypoxia by maximizing brain perfusion and thereby the supply of O_2_. Throughout this study, the effects of hypercapnia on brain and body *P*_O2_ were not due to an altered respiratory rate. There was either no observable change, or the rate decreased when P6 or P11 rat pups were exposed to 5%, 10% or 20% CO_2_.

Hypoxia alone (5% O_2_ in the inhaled gas) caused brain *P*_O2_ to fall to apparent zero (1.0 ± 0.5 mmHg, n = 4; Figure 2C) that was indistinguishable from the apparent zero level of brain *P*_O2_ seen during asphyxia (2.4 ± 0.9 mmHg, n = 8, *P* = 0.25). Recovery in air was associated with a rise above baseline in brain *P*_O2_ that developed rapidly like after asphyxia, but was much lower in amplitude (peak 45.6 ± 5.6 mmHg at 4.8 ± 0.6 min) and had a much longer duration (time from the beginning of recovery until mean *P*_O2_ fell below baseline: 49.9 ± 9.6 min after hypoxia vs. 19.9 min ± 4.9 min after asphyxia). The difference in brain *P*_O2_ recovery kinetics bears similarity with the slower brain pH recovery after hypoxia compared to asphyxia (Figures 2A,C vs. Figures 1A,C). Interestingly, a fall in brain *P*_O2_ to almost zero (2.6 ± 0.9 mmHg, n = 8) was observed also upon a moderate hypoxia (9% O_2_ in the inhaled gas; Figure 2D).

Taken together, the results illustrated in Figure 2 suggest, albeit do not demonstrate yet (see below), that CO_2_ can play an essential role in the control of neonatal brain tissue oxygenation during periods of compromised O_2_ availability. Our data indicate that brain consumes all available O_2_ during 5% O_2_ asphyxia and during 5% to 9% O_2_ hypoxia but, as such, a near-zero O_2_ level within brain parenchyma does not justify the conclusion that brain energy metabolism is primarily anaerobic under these conditions.

### 2.4 Intermittent asphyxia reveals an enhancing effect of elevated CO_2_ on brain oxygenation in P6 and P11 rats

In order to gain a deeper insight into the dependence of pH and *P*_O2_ on the developmental stage and severity of asphyxia, we used an experimental paradigm to mimic the intermittent O_2_ supply that is typical to BA.^26^ Here, we used both P6 and P11 pups which roughly correspond to preterm and full term babies in the developmental stage of the cortex.^19, 51^

In P6 rats, 9% / 5% O_2_ intermittent asphyxia (see scheme in Figure 3; and Materials and Methods) caused an acid shift and recovery in both brain and body pH with characteristics very similar to those seen in steady 5% O_2_ asphyxia, except for the smaller amplitude of acidosis in both compartments (brain 6.81 ± 0.01 and body 6.76 ± 0.014, n = 8, at the end of asphyxia; Figure 3A). In line with this, the BBB potential response differed from steady asphyxia mainly by its somewhat smaller amplitude. The alternation between 5% and 9% O_2_ in inhaled gas mixture gave rise to relatively small shifts (< 0.035) in both pH signals. The intermittent asphyxia is expected to induce a smaller metabolic acidosis than the steady one and this is, indeed, evident in the brain and body pH traces which diverge much less in the former (compare the shaded areas under ΔpH traces in Figures 1A and 3A).

**FIGURE 3.**
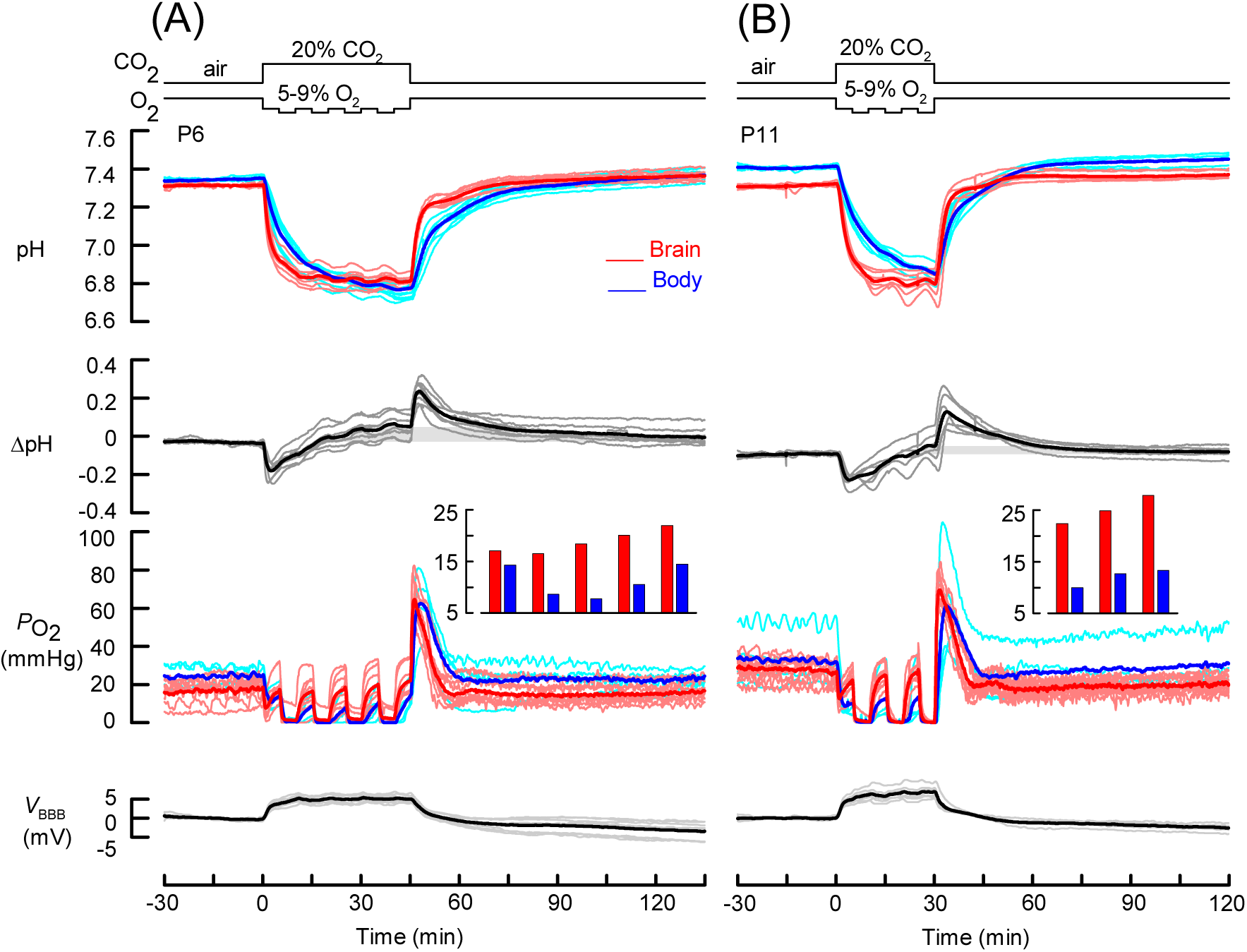
Intermittent asphyxia induced pH and *P*_O2_ responses in P6 and P11 rats. **(A)** Simultaneously recorded responses to 45 min intermittent asphyxia in brain and body pH as well as in *V*_BBB_ are shown in parallel with brain and body *P*_O2_ responses obtained in a separate series of otherwise identical experiments. All data in (A) are from P6 rats. **(B)** Similar to (A) but with a 30 min intermittent asphyxia applied on P11 rats. The bar graph insets in (A) and (B) show the maximum values of mean brain and body *P*_O2_ that were reached by the end of each 5 min period of inhaling 9% O_2_, 20% CO_2_.

An interesting observation on *P*_O2_ responses, particularly those in the brain, was made with intermittent asphyxia (n = 8 and 4 for brain and body, respectively; Figure 3A). At the beginning of asphyxia, in 9% O_2_/20% CO_2_, the two *P*_O2_ signals fell rapidly but started to recover within a minute indicating the activation of some compensatory mechanism(s). During the subsequent 5 min period with 5% O_2_, both brain and body *P*_O2_ fell to zero, in a manner similar to under conditions of 5% O_2_ asphyxia. When the O_2_ was then increased from 5% to 9% for 5 min, brain *P*_O2_ started to rapidly recover towards its control level, despite the continuous hypoxic level of ambient O_2_. During the subsequent three steps to the 9% level, the rise in brain *P*_O2_ became even larger, reaching – and in 5 of the 8 animals crossing – the pre-asphyxia control level during the last 9% O_2_ period (Figure 3A). Notably, the parallel partial recovery of body *P*_O2_ was much smaller, with its final peak at around half of the control level, indicating that the compensatory mechanism^15^ postulated above works in a more efficient manner in the brain.

Return to air evoked a transient overshoot in both brain and body *P*_O2_, very similar to those seen after 5% O_2_ steady asphyxia. Comparing to the steady asphyxia, the less severe nature of the intermittent asphyxia likely accounts for the lack of the body *P*_O2_ undershoot below its baseline level during the time interval of approximately 30 to 60 min after return to breathing air.

The above results clearly show that even a limited source of O_2_ during asphyxia can be sufficient for restoring normoxic conditions in the brain at P6. During development, increasing energy metabolism paralleled by angiogenesis^52, 53^ is expected to have consequences on brain pH and *P*_O2_ responses. Therefore, we next used the intermittent asphyxia model with P11 rats and, based on pilot experiments, we decreased the duration of exposure to 30 min to keep mortality at zero. The mean baseline brain pH of all P11 rats (7.31 ± 0.01, n = 11) did not differ from that at P6 (*P* = 0.91), but baseline body pH was slightly higher (7.41 ± 0.02, n = 11; *P* = 0.0043). The mean baseline brain *P*_O2_ in the P11 rats (31.7 ± 0.91 mmHg, n = 32), was noticeably higher than in P6 rats (*P* = 5 · 10^-12^). A comparable increase at P11 was seen in the body *P*_O2_ (34.2 ± 2.06 mmHg, n = 22; *P* = 0.0043). As is evident from Figure 3B, there were no obvious differences in the responses evoked by intermittent asphyxia at P11 compared to those at P6. In a series of experiments with hypercapnia like the one in Figure 2B but at P11 (not illustrated), the increases in *P*_O2_ upon any of the three CO_2_ levels (5%, 10%, 20%) differed neither in the brain nor in the body from those at P6 (*P* values from 0.38 to 0.94, n values from 4 to 10). Like in P6 rats, brain and body *P*_O2_ fell rapidly to zero during 9 % O_2_ hypoxia in P11 rats (15 min exposure; n = 6 and 4 for brain and body, respectively; not illustrated), and no obvious differences were seen in the *P*_O2_ responses during recovery compared to P6.

### 2.5 Neonatal guinea pigs maintain higher *P*_O2_ during severe asphyxia than rats

The guinea pig is a rodent which has been used in a number of translational studies on BA and other early-life disorders.^54^ This species is adapted to life at high altitudes^55, 56^ and it has precocial neonates, and the developmental stage of their cortex is much more advanced than in the altricial neonatal rat or even a full-term human newborn.^25^ Thus, to study further the developmental and inter-species aspects of brain *P*_O2_ responses, we used P6 and P11 rats as well as guinea pigs at P0–2 and compared their brain and body *P*_O2_ responses when exposed to steady 5% or 9% O_2_ asphyxia for 15 min.

As expected, 5% O_2_ asphyxia resulted in both P6 and P11 rats in a prompt fall in brain *P*_O2_ to apparent zero level, with little further change during the 15 min asphyxia (n = 10 and 7, respectively; Figure 4A top and middle panels). The recovery consisted in both age groups of a transient rise that peaked at a level that was approximately twice higher than the baseline *P*_O2_ value, followed by a slower fall to baseline level in about 15 min. The simultaneous body *P*_O2_ responses resembled those in the brain but had somewhat slower kinetics and lower recovery overshoot (n = 4 for both P6 and P11 rats).

**FIGURE 4.**
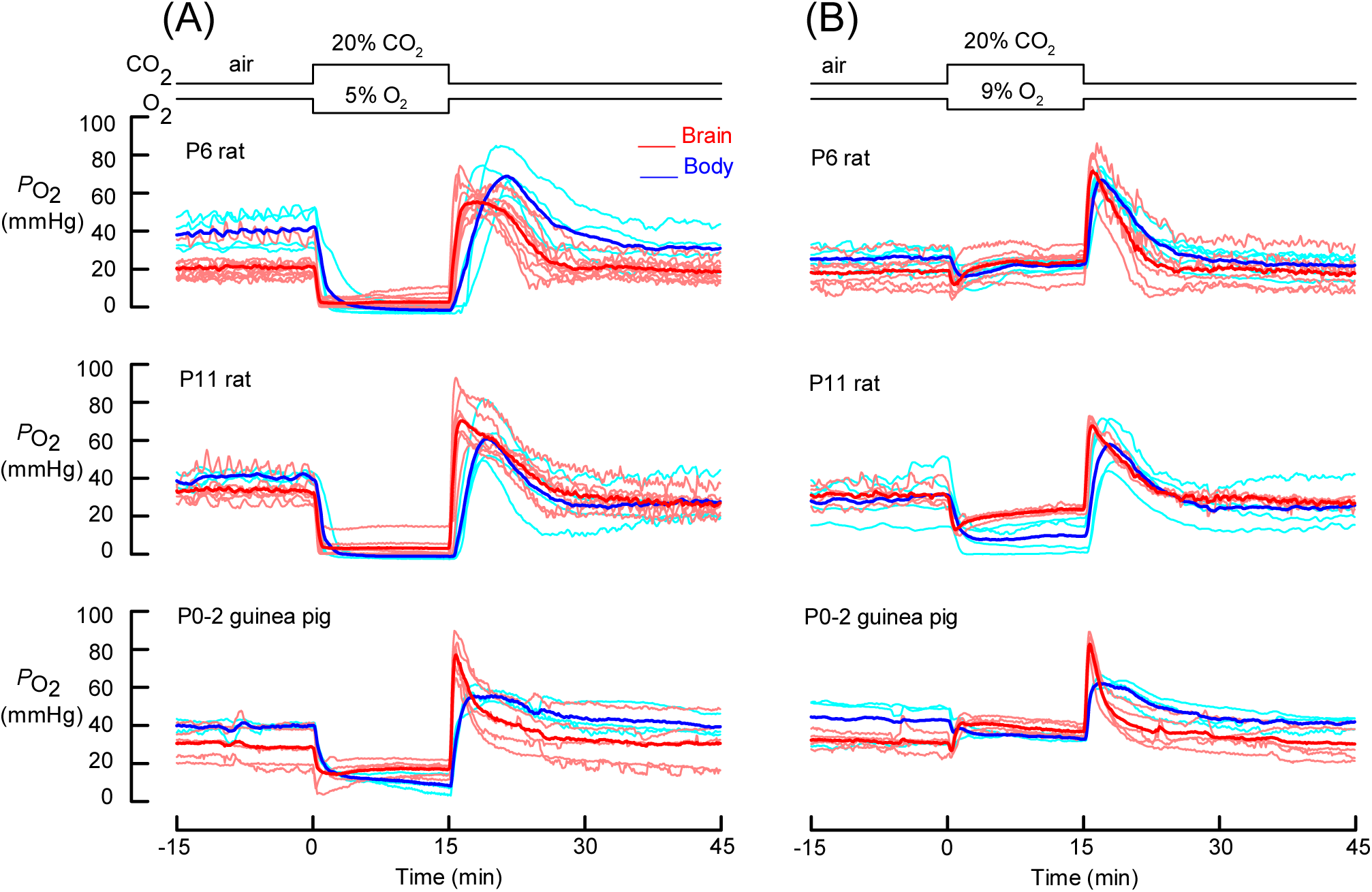
Brain and body *P*_O2_ responses to severe and moderate asphyxia in rats at P6 and P11 as well as in P0-2 guinea pigs. **(A)** Top, middle and bottom panels show superimposed individual and mean traces of brain and body *P*_O2_ recorded in P6 and P11 rats and P0-2 guinea pigs exposed to 5% O_2_ asphyxia for 15 min. **(B)** Similar to (A) but with a more moderate, 9% O_2_ asphyxia.

The moderate 9% O_2_ steady asphyxia for 15 min showed again recovery with the typical transient overshoot in P6 and P11 rats, but a small difference was observed between brain and body *P*_O2_ levels during the asphyxia period (Figure 4B top and middle panels). At P6, a very brief drop was followed by a rise in brain *P*_O2_ to a level that was at or even above the baseline whereas the body *P*_O2_ signal recovered from the initial drop more slowly and levelled off below its baseline (n = 6 and 5, respectively), i.e. brain and body were slightly hyperoxic and hypoxic, respectively, during the 9% O_2_ steady asphyxia. At P11, the initial drop in both brain and body *P*_O2_ was larger and only brain *P*_O2_ showed a partial recovery during asphyxia, reaching 76.7 ± 6.5% of its baseline value, whereas body *P*_O2_ levelled off at 27.1 ± 10.7% of its baseline level (n = 4 for both; Figure 4B middle panel).

In P0-2 guinea pigs, the baseline brain and body *P*_O2_ were 31.6 ± 1.6 mmHg (n = 5) and 43.2 ± 6.8 mmHg (n = 3), respectively. In a manner similar to P6 rats, guinea pig brain tissue did not become hypoxic during the 9% O_2_ asphyxia except for the very first <1 min, but instead rapidly increased to a level that was 9.2 ± 2.0 mmHg above baseline at 5 min, and then slowly declined towards the baseline level before a very brief overshoot to 82.9 ± 2.3 mmHg was evoked by return to breathing air (n = 5; Figure 4B bottom panel). The similarity to P6 rats was seen also in the response of guinea pig body *P*_O2_ to 9% O_2_ asphyxia (by the end of asphyxia, body *P*_O2_ fell to 33.1 ± 0.7 mmHg, n = 3, i.e. to 81.3 ± 15.4% of baseline). Interestingly, during the severe 5% O_2_ asphyxia, guinea pigs differed from both P6 and P11 rats in that their brain *P*_O2_ fell rapidly to a transient minimum of 35.4 ± 5.4% of the pre-asphyxia baseline and then increased and stayed at no less than 61.0 ± 9.9% of baseline (16.9 ± 1.9 mmHg) till the end of asphyxia (n = 5; Figure 4A bottom panel). The simultaneously recorded body *P*_O2_ showed a more robust, progressive fall that reached 8.7 ± 3.2 mmHg, i.e. 22.0 ± 8.1% (n = 3) of the pre-asphyxia baseline, by the end of the 15 min 5% O_2_ asphyxia, followed by a moderate overshoot during recovery.

### 2.6 Effects of GRN on brain and body pH, and on brain *P*_O2_, during recovery from asphyxia in P6 and P11 rats

GRN applied during recovery from asphyxia holds promise as a therapeutic intervention that protects the brain and improves the outcome.^39^ Therefore, in an extensive series of experiments we focused on steady and intermittent asphyxia with GRN.

In contrast to the fast pH recovery seen in P6 rats during rapid restoration of normocapnia (RRN) after 45 min steady 5% O_2_ asphyxia (Figure 1A), the recovery that was seen during GRN had three distinct phases in both brain and body (Figure 5A). As expected, these phases in pH recovery paralleled those in the ambient CO_2_ levels, and therefore the recovery of both brain and body pH to baseline was much slower than with RRN. During GRN, body pH remained below brain pH (see ΔpH in Figure 5A), and the final pH levels at the end of recovery were identical with those seen with RRN (*P* = 0.62 and *P* = 0.76, respectively). Again, the *V*_BBB_ signal closely followed the time course of pH changes. Sudden *V*_BBB_ collapses were not associated with any of the experimental insults used in this study, which suggests that the insults did not cause BBB disruption.

**FIGURE 5.**
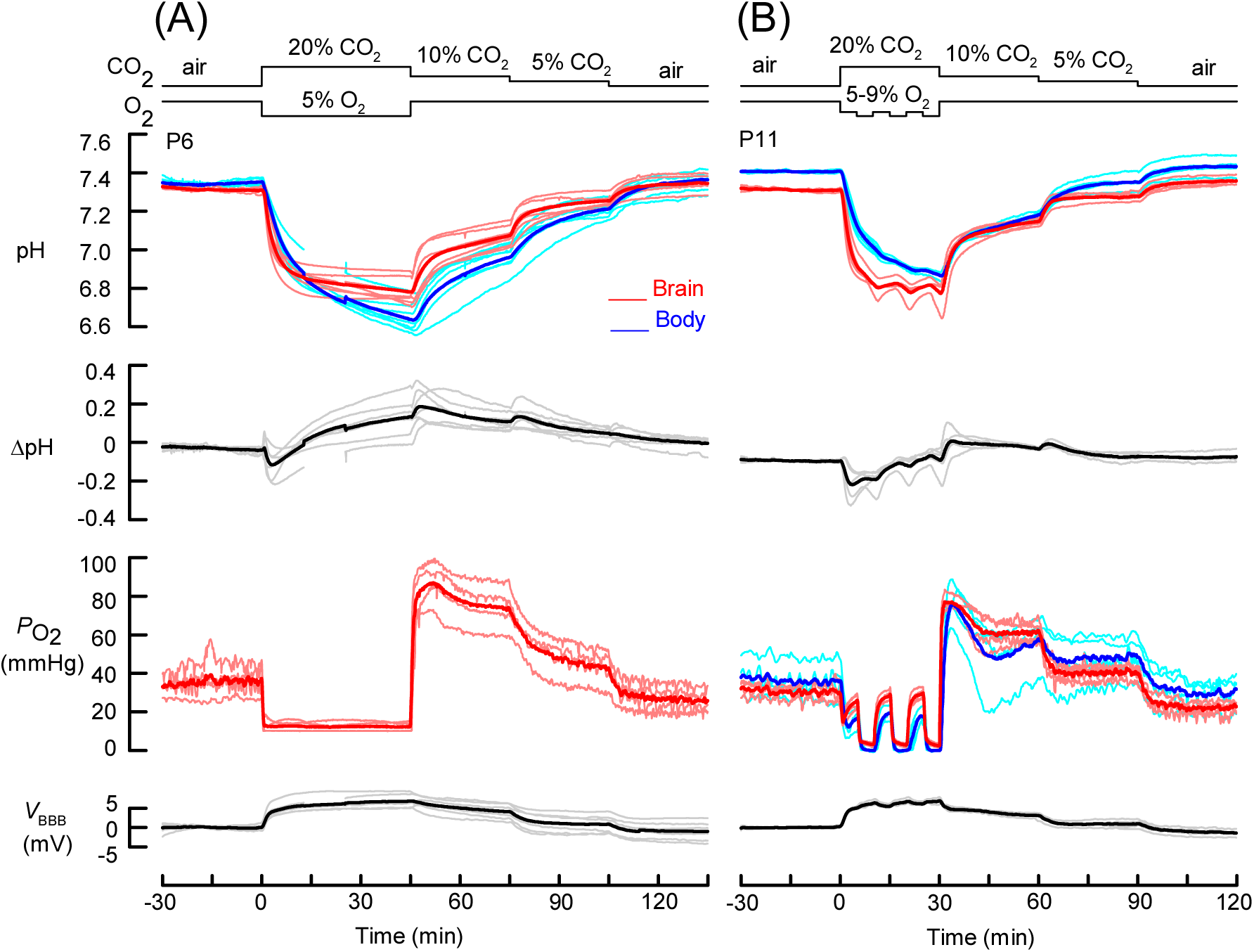
Graded restoration of normocapnia during recovery from asphyxia slows down the alkaline recovery and prolongs the *P*_O2_ overshoot in P6 and P11 rats. **(A)** P6 rats were exposed to 5% asphyxia for 45 min after which recovery took place with graded restoration of normocapnia, i.e., normal ambient O_2_ level was restored immediately whereas CO_2_ in the inhaled gas was first decreased to 10% for 30 min followed by 5% CO_2_ for another 30 min before the animals were finally breathing normal air. Superimposed individual and mean traces of simultaneously recorded brain and body pH and *V*_BBB_ are shown in parallel with brain *P*_O2_ data from a separate series of otherwise identical experiments. **(B)** Similar to (A) but the experiments were done on P11 rats and using intermittent asphyxia for 30 min, and also body *P*_O2_ was recorded simultaneously with brain *P*_O2_. One of the body *P*_O2_ traces deviates from corresponding traces recorded from other P11 rats and could therefore be considered as an outlier in data. However, since we were unable to identify any technical reasons for this atypical behaviour during early recovery, we did not discard this recording from the data analysis.

In parallel experiments at P6, the transient post-asphyxia overshoot in brain *P*_O2_ peaked rapidly at a level (77.9 ± 5.7 mmHg, n = 4) that was nearly three times higher than the baseline, and higher and broader than the peak seen during RRN (cf. Figure 2A; *P* = 0.31 for peak height comparison). Thereafter, *P*_O2_ decreased slightly (to 63.6 ± 6.1 mmHg) by the end of the 30 min exposure to 10% CO_2_ / 20% O_2_ gas, followed by a further fall in brain *P*_O2_ when CO_2_ was lowered to 5%. The fall was slow and did not attain a stable level in 30 min (brain *P*_O2_ 60 min after end of asphyxia 32.5 ± 5.0 mmHg). Return to air 60 min after the end of asphyxia made brain *P*_O2_ fall to a somewhat hypoxic level.

The effects of GRN with P11 rats after 30 min intermittent asphyxia (n = 4; Figure 5B) were consistent with those described above for P6 rats. The peak in brain *P*_O2_ was higher and broader than with RRN at P11. Compared to GRN at P6, a stable brain *P*_O2_ level was attained more rapidly during the 30 min recovery periods with 10% and 5% CO_2_, and these levels were approximately 2 and 1.3 times higher than the mean baseline, respectively. Return to air caused a fall in brain *P*_O2_ to a hypoxic level (21.5 ± 1.3 mmHg) followed by a slow trend towards the original baseline value.

## 3 DISCUSSION

In this study we have focused on pH and *P*_O2_ as two key variables in the brain *milieu interior* in our rodent model of BA. We have examined the magnitude and time course of the perturbations caused by a number of experimental paradigms, including *severe* and *moderate asphyxia*, as well as severe and moderate *hypoxia* and *hypercapnia* in isolation (see Materials and Methods). This permitted the identification of adaptive mechanisms pointing to the remarkable ability of the immature rodent brain to cope with the low O_2_ levels associated with asphyxia, as has been previously shown in extensive studies on larger mammals, such as sheep and pigs.^14–16^ Moreover, our experiments indicate that these adaptive mechanisms are not at work during pure hypoxia, which is a condition that a mammal never faces – pre-, peri- or postnatally – under natural conditions. The present data will also increase our understanding on the conditions *in vivo* which will affect neuronal excitability during BA. Finally, we demonstrate that *graded restoration of normocapnia* after a period of asphyxia will prolong the actions of the innate brain-protective mechanisms as well as those of hypercapnia itself.

<u>Our main observations on pH</u> are that (i) while the hypercapnia component of asphyxia is responsible for most of the large and fast acidosis seen in brain and body pH, the brain appears to be protected against the more slowly developing metabolic acidosis. We found that (ii) unlike asphyxia, exposure to hypoxia produces a small but immediately triggered brain-confined *alkalosis* which is paralleled by a slow and similarly modest (by definition a metabolic) acidosis in the body, pointing to the BBB’s capability of acid extrusion^57^ at a rate that exceeds net acid production in the brain, as postulated earlier on,^58^ see also ref. [^59^]. The hypoxia-induced net brain alkalosis shows that BA cannot be adequately mimicked by *in vivo* hypoxia (see below). Finally, (iii) the fast rate of recovery of brain pH after asphyxia was significantly reduced by GRN, an effect that has a pronounced suppressing action on neocortical seizures as shown by Ala-Kurikka *et al.*^60^ in a parallel study. Notably, in the present work done with anaesthesia and a slightly lower body temperature of 33.5 – 34 °C vs. 36.5 – 37 °C, no seizures took place.

<u>With regard to oxygen levels</u>, we found a striking difference if hypoxia was applied alone, or whether it was applied as one of the two components of asphyxia. During hypoxia (5% or 9% O_2_) as well as during severe experimental asphyxia (i.e., 20% CO_2_ plus 5% O_2_), both brain and body *P*_O2_ fell to levels that were close to zero. However, with 9% O_2_ in parallel with 20% CO_2_ hypercapnia (moderate asphyxia), the brain regained its control *P*_O2_ level pointing to the activation of highly effective adaptive mechanisms. Moreover, our recordings show a prompt post-asphyxia overshoot of brain (and body) *P*_O2_ which was prolonged by GRN.

The above effects as well as their mechanisms and consequences will be discussed in detail below.

### 3.1 Towards a valid rodent model of birth asphyxia

#### Characteristics of the present model

Choosing an experimental model to study BA is not a straightforward task. Obviously, *in vitro* models cannot reproduce systems level-adaptive responses to asphyxia. Because of the altricial nature of rat and mouse offspring, their cortical development corresponds around P6 to human preterm (from 28 up to 35 gestational weeks) and at around P11 to term babies.^19, 21, 22, 51^ Therefore, experimental manipulations must be designed in a manner that reproduces asphyxia-mimicking conditions in these postnatal animals, which have already acquired a fully functional pulmonary ventilation.

The main features of our model are the following: (i) It is noninvasive, and therefore the endogenous vasomotor and other systems-level^61, 62^ (e.g. neurohormonal)^24, 47^ protective mechanisms remain fully operational. ii) Compromised gas exchange via the umbilical cord, which results in both CO_2_ accumulation and O_2_ deficit in the fetus, is mimicked by elevating CO_2_ and reducing O_2_ in the ambient gas, instead of exposing the animal to a hypoxic gas which blocks the respiratory acidosis characteristic of asphyxia (see section on Asphyxia vs. hypoxia below). iii) The experimental manipulations target the entire infant rat, with a tissue and organ distribution of adaptive and other physiological responses which are of endogenous origin. iv) Furthermore, our pH and lactate data indicate that the current model reproduces the key clinical criteria of severe BA in human neonates, namely systemic acidosis to pH levels <7.0 and a base deficit ≥12 mmol L^-1^.^8, 63, 64^ Regarding hypercapnia, umbilical cord artery gas partial pressure (*Pa*) values in neonates with severe acidosis at birth are typically >100 mmHg *Pa*_CO2_ (≥140 mmHg in >10% of cases), and *Pa*_O2_ is often in the range 10 to 15 mmHg.^65, 66^ Thus, in all the three versions of the present model (moderate, severe and intermittent asphyxia), CO_2_ was maintained at the constant 20% level, while O_2_ was reduced to 9% or 5% in order to unravel the animal’s fully activated endogenous capacity to maintain brain oxygenation despite the limited oxygen resources.

The newborn guinea pig is precocial and therefore its brain at birth is much more mature than that of the altricial rat.^51, 52, 67^ However, our results with P0 to P2 guinea pigs (which correspond to two to three week old rats in terms of cortical development)^25, 52^ were qualitatively identical to, and even quantitatively similar, to the results obtained with the P6 and P11 rats, suggesting that the present experimental approaches are valid in translational work on all standard laboratory rodents.

#### Asphyxia vs. hypoxia

Our present data on brain alkalosis induced by pure hypoxia speaks against the relevance of hypoxia models of asphyxia. Another important line of evidence comes from experiments on hypoxia on neuronal excitability. Despite a number of purinergic mechanisms activated during oxygen deprivation,^68, 69^ *in vivo* hypoxia alone is known to produce a gross increase in cortical excitability which becomes manifest as seizures *during* this challenge.^32–35^ If, indeed, the hypoxia models are intended to mimic what happens in clinical BA, this would imply that the neonatal seizures associated with complicated birth are triggered already *in utero* and during parturition. This is obviously not the case. What is missing in hypoxia models of BA is hypercapnia, and the consequent respiratory acidosis that lowers brain excitability by modulating a wide variety of voltage and ligand-gated ion channels, see ref. [^29^] and references cited therein.

Thus, while hypoxia promotes neocortical excitability, hypercapnic acidosis has a functionally opposite effect. This is, notably, consistent with the fact that neonatal seizures caused by asphyxia take place with a substantial delay (several hours) after birth,^70^ i.e. at a time when normoxic conditions have already been established. From a general neurobiological point of view, it is interesting to note that, under all natural conditions, hypoxia *in vivo* is associated with hypercapnia, suggesting an evolutionary history of the development of the neuromodulatory effects of pH *in vivo*,^29^ whereby neuronal acidosis suppresses excitability.^71–73^

Thus, the recovery of brain pH is likely to be a factor that sets the time course of the increase in brain excitability during recovery from asphyxia. Worth mentioning is that in our previous study^39^ we used isoflurane instead of urethane anaesthesia, and the initial depth of anaesthesia may have been too deep, likely accounting also the significantly (by >0.1 units) lower baseline brain pH compared to the present study. These differences may explain why in the present study we did not detect a net post-asphyxia brain alkalosis as large as the one described in previous work.

#### Brain-protecting systems-level mechanisms

To spare the fetal brain which is critically dependent on oxygenation, asphyxia triggers systems-level endogenous protective mechanisms which are usually referred to as the peripheral chemoreflex or the brain sparing effect. These protective responses involve vasodilation and vasoconstriction which act to maintain perfusion of vital, highly oxygen-dependent organs like the brain, heart and adrenal glands,^13–15, 74^ thereby reducing the risk of HIE following BA.^40^

While studies monitoring CBF have shed light on adaptive responses acting to maintain brain oxygenation, it is obvious that CBF data do not provide direct information on the level oxygen in brain tissue during asphyxia. For instance, during asphyxia, CBF increases but at the same time there is not only a large decline in O_2_ availability but also a large fall in blood pH, which reduces the oxygen carrying capacity of blood because of the Bohr effect.^75–77^ Thus, measurements of *P*_O2_ in brain tissue provide direct quantitative information on this key variable, as is the case in the present study. A direct demonstration of the CO_2_-dependence of brain oxygenation during asphyxia is that while hypoxia brought about by lowering ambient O_2_ to 9% causes a near-instantaneous fall in brain *P*_O2_ to about one tenth of the baseline level, nearly normoxic conditions prevail in the brain parenchyma when the same level of hypoxia is combined with hypercapnia. The brain *P*_O2_ responses to 9% O_2_ during intermittent asphyxia appeared as if this brain sparing effect is augmented by the steps to the low (5%) O_2_ levels – a finding similar to what has previously been reported in response to umbilical cord occlusions in a sheep model of BA.^78^ However, the increasing trend in brain *P*_O2_ as observed in the peak levels reached by the end of each 9% O_2_ period (see Figure 3 bar graph insets) does not differ from that seen during steady 9% O_2_ asphyxia, indicating a similar augmentation of the chemoreflex during steady and intermittent asphyxia in the present experimental setting.

Our finding that not only brain *P*_O2_ but also body *P*_O2_ (measured subcutaneously) increased upon hypercapnia does not imply that the two are based on identical mechanisms. Subcutaneous *P*_O2_ increases upon hypercapnia also in humans, but this effect is caused by an increase in cardiac output,^79, 80^ and not by vasodilation, which is known to be the key mediator of the increase in CBF during hypercapnia.^13^

The results obtained with P6 and P11 rats showed only minor differences between these two developmental stages. Thus, as far as systems-level metabolic and brain-sparing aspects of asphyxia are considered, our model can be used to study perinatal asphyxia at different stages of cortical development, corresponding to preterm to term human babies. Inducing asphyxia via the inhaled gas provides a straightforward method to generate both steady and intermittent asphyxia protocols. These protocols correspond, in rough terms, to two mechanistically different perinatal complications: to acute placental insufficiency and to prolonged periods of uterine contractions, respectively.

### 3.2 Graded restoration of normocapnia

As discussed above (see Results), the acidosis during experimental asphyxia had two distinct components, respiratory and metabolic, whereof the former is larger in amplitude in both brain and body, and caused by hypercapnia. GRN slowed down the pH recovery after asphyxia by prolonging the duration of respiratory acidosis, and prolonged the overshoot in brain and body *P*_O2_. Whether these effects are beneficial with regard to outcome following asphyxia is not immediately obvious. However, a post-asphyxia seizure-suppressing action of lower brain pH during GRN can be readily assumed (see above, and e.g. ref. [^73^]) and this kind of an effect has been directly demonstrated in a parallel study by Ala-Kurikka *et al.*^60^

By definition, hypercapnia causes acidosis without causing any base deficit; and the generation of the base deficit is strictly dependent on metabolic (lactic) acidosis.^81, 82^ This implies that the lower pH maintained by GRN during recovery from asphyxia does not enhance the base deficit. It may be speculated that since acidosis, no matter if metabolic or respiratory, can limit lactic acid generation,^83^ GRN may even assist in the restoration of normal base content.

Spontaneous hypocapnia resulting from hyperventilation is not uncommon in asphyxiated term babies, and hypocapnia occurs often in ventilated – especially preterm – infants. An association between hypocapnia and adverse outcome has been shown in clinical studies on neonates with HIE.^84^ A neuroprotective effect of CO_2_ against hypoxic-ischemic injury has been demonstrated in rats during hypoxia^85^ which, in fact, merely converts the experimental insult from hypoxia to asphyxia. Hypocapnia is particularly injurious to the preterm human brain during the first days of life as it causes brain hypoperfusion and often leads to development of periventricular leukomalacia.^86, 87^ Mild permissive hypercapnia has been repeatedly suggested as a safe manipulation that can reduce lung injury due to bronchopulmonary dysplasia in ventilated preterm neonates, however no general recommendations for its optimal use have existed and the levels of hypercapnia with beneficial rather than adverse consequences remain unclear.^88–90^ Regarding the current situation with contradicting studies on this issue, it is worth pointing out that the above studies on permissive hypercapnia focus on blood gases in human newborns over a time period of several days or weeks after birth. For instance, in a study on ventilated very low birth weight preterm infants, hypercapnia during the first week of life was found to lead to a progressive loss of cerebral autoregulation with an associated risk of brain injury.^91^ In contrast, GRN as studied here, is applied immediately after asphyxia, corresponding to the first hour of extrauterine life after a complicated birth. This is the time when severely asphyxiated newborns often have unstable blood gas levels with episodes of severe hyperoxaemia and severe hypocapnia (arterial *P*_O2_ and *P*_CO2_ >200 mmHg and <20 mmHg, respectively), where these fluctuations *per se* are thought to have a major contribution to brain injury.^92^ Given the steep dependence of *P*_O2_ on *P*_CO2_, a strategy to maintaining stability in *P*_O2_ might be based on permissive hypercapnia or on a modification of GRN.

The transient overshoot that was seen in brain and body *P*_O2_ in all our experiments during the first 15 minutes of recovery from asphyxia with RRN is readily accounted for by normal air becoming suddenly available when the endogenous compensatory mechanisms triggered by asphyxia are still acting. In addition to increased CBF, inhibition of mitochondrial respiration and reduced oxygen consumption are likely to be involved.^93–95^ The transient *P*_O2_ overshoot is most likely exaggerated in P6 or P11 rats that have acquired fully functional pulmonary gas exchange unlike human newborns, and therefore the amplitude and duration of the *P*_O2_ transient during early recovery should be considered within the framework of the model. Notably, in experiments with RRN, the overshoot was followed by a tendency towards hypoxic *P*_O2_ levels for over an hour in experiments with RRN, whereas elevated *P*_O2_ levels were seen in both brain and body as long as the inhaled gas contained CO_2_ in the GRN experiments. These data point to a beneficial action of GRN on brain oxygenation during the onset of pulmonary ventilation.

The brain hyperoxia seen during GRN might raise concerns about its neurotoxic effects. However, unlike with hyperoxic resuscitation^96^ the fraction of inspired O_2_ is not elevated during GRN, and the *P*_O2_ peak associated with GRN is briefer than what is typically used in neonate animal models of oxygen toxicity.^97^ More importantly, hyperoxia occurs in parallel with the GRN-induced hypercapnic acidosis. Therapeutic hypercapnia during reperfusion has been shown to attenuate inflammation and to reduce free radical-mediated injury in an *in vivo* rabbit model of ischemic lung injury,^98, 99^ and slowing down the abrupt increase in pH during reoxygenation was found to reduce anoxic injury in perfused rat livers.^100^ Based on their own and others’ data Halestrap and coworkers conclude that significant superoxide production in the mitochondrial matrix is unlikely during the first minutes of reperfusion in cardiac cells, and that a large increase in intracellular reactive oxygen species occurs only after opening of mitochondrial permeability transition pores,^101, 102^ which remain inhibited at pH values below 7. Thus, there are reasons to think that the short-term hyperoxia that is linked to the GRN-based hypercapnic acidosis is not detrimental as such.

Taken together, in the present work, the rationale of using GRN as a brain protecting intervention during early recovery from asphyxia is based on augmenting endogenous neuroprotection^40^ by making the activity of the brain sparing mechanisms outlast the asphyxia period. Here, we want to emphasize that using the present GRN protocol is a translational proof-of-concept study, where the amplitudes and durations of the descending CO_2_ levels (10% and 5% CO_2_) are not intended to be tested as such in the clinic. In fact, we have convincing preliminary evidence^60^ that a much milder GRN protocol based on 5% CO_2_ provided post-asphyxia leads to a suppression of neocortical seizures following the intermittent asphyxia paradigm in P11 rats.

### 3.3 The blood-brain barrier

*V*_BBB_ is a transendothelial potential difference that prevails between brain tissue and blood. Respiratory pH changes cause large shifts in *V*_BBB_ that have been measured in invasive recordings in experimental animals and at scalp in human subjects.^48, 49^ In the present experiments the reference electrodes of the brain and body pH microelectrodes measured tissue potentials, and *V*_BBB_ was obtained as their difference. *V*_BBB_ responses to asphyxia were smooth and they appeared to follow primarily the changes in body pH (≈ blood pH) – this was evident during hypoxia when brain pH and body pH shifted in opposite directions. However, there are no grounds to assume that *V*_BBB_ depends only on one variable, since it originates as the difference of the apical and basolateral membrane potentials of the BBB forming cells. If a large-scale BBB disruption occurs, it shunts *V*_BBB_ and a transient shift is seen in the *V*_BBB_ signal.^103, 104^ We did not find any indication of robust BBB disruption upon asphyxia in the present experiments. Instead, the present results suggest that BBB remained at least largely if not fully intact.

The fact that *V*_BBB_ can be easily measured noninvasively^49, 104^ raises the idea that using one EEG channel dedicated to the monitoring of DC shifts may open up a new window for brain monitoring following BA.

## 4 MATERIALS AND METHODS

### 4.1 Ethical Approval

All experimental procedures were carried out in accordance with the European Convention for the Protection of Vertebrate Animals used for Experimental and Other Scientific Purposes (ETS 123) and the Directive 2010/63/EU, implemented in Finland in Act 497/2013 and Decree 564/2013 on the protection of animals used for scientific or educational purposes. All experiments were approved by the Local Animal Ethics Committee of Helsinki University and the National Animal Ethics Committee in Finland.

### 4.2 Animals

Experiments were performed on male Wistar rats on P6 (*n* = 96) or P11 (*n* = 43), or on P0 to P2 guinea pigs of either sex (*n* = 5). Since none of the responses was tested multiple times using the same animal, *n* indicates the number of experiments as well as animals throughout the paper. Rats were obtained immediately before experiments from an in-house animal facility operating under control of the Laboratory Animal Centre of the University of Helsinki (LAC), where rats were housed in cages under 12-hour light/dark cycle and with access to food and water *ad libitum*. Guinea pigs were maintained under similar conditions in a LAC facility from where they were obtained no longer than 30 min before they were anaesthetized for surgery.

### 4.3 Surgery and preparation for the recordings

Animals were anaesthetized with 4% isoflurane in room air, 1 mg g^-1^ urethane was given by intraperitoneal injection and isoflurane was reduced to 1.5 – 2% for surgery. After removal of the scalp skin and soft tissue, craniotomies were made for the electrode implantation, including: (1) craniotomies over the hippocampi for the pH (two 0.9 mm holes contralateral to each other) and *P*_O2_ (one 0.9 mm hole) recordings; (2) craniotomy in the occipital bone for the ground wire (placed over the cerebellum). The exposed dura was gently opened using a fine needle to allow electrode implantation. Skin incisions were made in the lower back for the subcutaneous placement of the body pH and *P*_O2_ recordings. At the end of surgery, an additional dose of 0.5 mg g^-1^ urethane was injected and isoflurane removed completely. If animal reacted to tail pinch during sensor placement, an additional dose of urethane (0.5 mg g^-1^) was injected before the beginning of baseline recording.

After surgery, the animal was placed on a warming pad in the experimental setup, with its head fixed to a stereotactic device. Body temperature was controlled with a rectal probe and BAT-12 thermometer (Physitemp, New Jersey, USA), and the heating was adjusted to maintain body temperature at the level of 33.5 to 34.0 °C during baseline recording in rats to avoid mortality during hypoxia and asphyxia.^39^ In the guinea pig experiments, heating settings were kept similar to those used with rats. A piezo movement sensor (PMS20S, Medifactory, Heerlen, The Netherlands) was attached on the lower part of chest with tape to record the respiratory rhythm.

### 4.4 pH and *P*_O2_ recording

Commercial H^+^-sensitive glass-membrane pH microelectrodes, models pH-25 and pH-500, as well as Clark-type polarographic O_2_ microsensors, models OX-10 and OX-N (Unisense A/S, Aarhus, Denmark) were used for pH and *P*_O2_ recordings in brain and body, respectively. Tip diameters of the sensors were: 10 μm for brain *P*_O2_, 25 μm for brain pH, 500-750 μm for body pH and *P*_O2_. Glass capillary micropipettes with tips broken to an approximate outer diameter of 20 μm and filled with 0.9% NaCl were used as reference electrodes for pH recording. Brain and body pH signals and their reference-electrode signals were recorded using custom-made electrometer amplifiers and an extracellular field potential amplifier (EXT-02F/2, npi electronic GmbH, Tamm, Germany). Brain and body *P*_O2_ signals were recorded using a PA2000 Picoammeter (Unisense A/S, Aarhus, Denmark). Signals were anti-alias filtered and digitized using Micro1401-3 converter (CED, Cambridge, UK), with sampling frequencies of 100 Hz for breathing and 10 Hz for pH, *P*_O2_ and temperature, and recorded on hard disk with Spike 2 software. The blood-brain barrier potential (*V*_BBB_) was measured as the difference between brain and body reference electrodes.

Coordinates for the brain probe implantations were 3-3.5 mm posterior, 3-4 mm lateral, 2.5-3 mm depth from bregma for the P6 and P11 rats, 5 mm posterior, 6 mm lateral, 3.5 mm depth from bregma in guinea pigs. The reference electrode for brain pH recording was always contralateral to the pH microelectrode, and ipsilateral to the brain *P*_O2_ microsensor in animals with simultaneous brain *P*_O2_ and pH recording. Body pH, *P*_O2_ and reference electrodes were placed subcutaneously, with tips advanced at least 10 mm from the skin incision. Skin incisions were covered with silicone grease to prevent air from accessing the site of recording. In contrast to other probes used in this study, body *P*_O2_ baseline values, unlike the responses to changes in experimental conditions, were slightly influenced by the positioning of the probe, resulting in a wider range of baseline values.

### 4.5 Calibration of pH and *P*_O2_ sensors

Calibration of all pH and *P*_O2_ sensors used was done before and after each experiment. pH electrodes were calibrated with their reference electrodes using two solutions containing 150 mmol L^-1^ NaCl and 20 mmol L^-1^ HEPES, pH adjusted to 6.8 and 7.8 with NaOH. The exact pH values of calibration solutions were regularly checked with a standard laboratory pH meter. O_2_ sensors were calibrated using standard extracellular solution ^24^ bubbled for at least 30 minutes with two gas mixtures, one containing 0% O_2_ (5% CO_2_ in N_2_) and the other containing 5 or 9% O_2_ and 5-20% CO_2_ in N_2_. All calibrations were done at room temperature, and a temperature correction was applied during data analysis.

Since pH calibrations were done at room temperature, the temperature sensitivity of the differential pH recording was found out experimentally, not forgetting the temperature dependence of the pKa of HEPES used in the calibration solutions. Based on the results, a correction of −0.09 pH units was applied to tissue pH values. Differences in observed brain or body pH baseline values between individuals in a cohort reflect true differences in pH as well as random sources of error characteristic of the method. The latter is likely to dominate, and therefore baselines of individual recordings were offset to the corresponding means.

The signal of polarographic O_2_ microsensors and the solubility of gaseous O_2_ are temperature dependent. Thus, in order to quantify the data in terms of partial pressure, we analysed the overall effect of these temperature dependencies using solutions that were vigorously bubbled for at least half an hour at room temperature (20-21 °C) and at 34 °C with gas mixtures containing 0%, 5% or 9% O_2_, and found a temperature dependence of ∼1% per °C. Brain and body *P*_O2_ sensor data were corrected accordingly during data analysis. At very low tissue *P*_O2_ values (sensor current close to 0 pA) that typically occurred during 5% hypoxia or asphyxia, the *P*_O2_ trace was still fluctuating or it was steady, showing no fluctuation for several minutes. In experiments where the latter condition occurred, the non-fluctuating level was taken as “true zero”, and the *P*_O2_ trace was offset accordingly. The average offsets of this kind were 1.5 mmHg and 2.5 mmHg for brain and body *P*_O2_, respectively, and applied to traces where a true zero condition was not seen during the experiment.

### 4.6 Experimental protocols

After initial stabilization of the recorded signals, a 30 min baseline was recorded from each animal breathing humidified room air applied via a small-rodent facemask (model OC-MFM for rats, OC-LFM for guinea pigs, World Precision Instruments, Sarasota, USA), at a flow rate of approximately 1200 ml min^-1^. The humidified experimental gas mixtures (AGA [Linde Group], Finland) were applied at the same rate, and were as follows: 5% or 9% O_2_ and 20% CO_2_ in N_2_ (asphyxia); 5% or 9% O_2_ in N_2_ (hypoxia); 5%, 10% or 20% CO_2_ and 20-21% O_2_ in N_2_ (hypercapnia; the first two also for GRN). For clarity, the timing of gas applications are given in the Figures using schematic traces.

### 4.7 Lactate measurement

Blood lactate was measured with a GEM Premier 4000 blood gas analyser (Instrumentation Laboratory, Bedford, MA, USA). P6 rats were exposed to 5% asphyxia for 15 min, immediately after which blood was collected and analysed as described before.^47^

### 4.8 Data processing and statistics

Data were processed using custom-made scripts in Matlab (MathWorks Inc.), and with Excel (Microsoft) and SigmaPlot 14 (Systat Software, Inc.).

All numerical data are given as mean ± SEM. Differences between mean values were assessed using unpaired or paired t-tests, and if not specified the test was unpaired. *P* < 0.05 was considered statistically significant. We are aware of the limitations of the t-test when sample sizes are small, and we give *P* values as suggestive information only. The reader might appreciate the low variability in the primary data, which is evident in the Results.

## ACKNOWLEDGEMENTS

We thank Maria Partanen, Ann-Christine Aho and Auli Kiukkonen for the maintaining and breeding of rats and guinea pigs. This work was supported by Grant ERC-2013-AdG 341116 (KK), Academy of Finland Grants 319237 and 294375 (KK) and the Emil Aaltonen Foundation Grant 180206 N1V (MP)

## AUTHOR CONTRIBUTIONS

KK and JV conceived the study and designed the experiments with AP and MP. The data were collected by AP and analysed by AP and JV. All authors were involved in data interpretation and writing the manuscript, and approved the final version of the manuscript.

## REFERENCES

1. Liu L, Oza S, Hogan D, Perin J, Rudan I, Lawn JE, Cousens S, Mathers C, Black RE. Global, regional, and national causes of child mortality in 2000-13, with projections to inform post-2015 priorities: an updated systematic analysis. Lancet. 2015;385:(9966):430–40.

2. Pisani F, Orsini M, Braibanti S, Copioli C, Sisti L, Turco EC. Development of epilepsy in newborns with moderate hypoxic-ischemic encephalopathy and neonatal seizures. Brain Dev. 2009;31:(1):64–8.

3. Ahearne CE, Boylan GB, Murray DM. Short and long term prognosis in perinatal asphyxia: An update. World J Clin Pediatr. 2016;5:(1):67–74.

4. Modabbernia A, Mollon J, Boffetta P, Reichenberg A. Impaired Gas Exchange at Birth and Risk of Intellectual Disability and Autism: A Meta-analysis. J Autism Dev Disord. 2016;46:(5):1847–59.

5. Hisle-Gorman E, Susi A, Stokes T, Gorman G, Erdie-Lalena C, Nylund CM. Prenatal, perinatal, and neonatal risk factors of autism spectrum disorder. Pediatr Res. 2018;84:(2):190–8.

6. Pappas A, Korzeniewski SJ. Long-Term Cognitive Outcomes of Birth Asphyxia and the Contribution of Identified Perinatal Asphyxia to Cerebral Palsy. Clin Perinatol. 2016;43:(3):559–72.

7. Gunn AJ, Bennet L. Brain cooling for preterm infants. Clin Perinatol. 2008;35:(4):735-vii.

8. Azzopardi DV, Strohm B, Edwards AD, Dyet L, Halliday HL, Juszczak E, Kapellou O, Levene M, Marlow N, Porter E, Thoresen M, Whitelaw A, Brocklehurst P. Moderate hypothermia to treat perinatal asphyxial encephalopathy. N Engl J Med. 2009;361:(14):1349–58.

9. Johnston MV, Fatemi A, Wilson MA, Northington F. Treatment advances in neonatal neuroprotection and neurointensive care. Lancet Neurol. 2011;10:(4):372–82.

10. Azzopardi D, Strohm B, Marlow N, Brocklehurst P, Deierl A, Eddama O, Goodwin J, Halliday HL, Juszczak E, Kapellou O, Levene M, Linsell L, Omar O, Thoresen M, Tusor N, Whitelaw A, Edwards AD. Effects of hypothermia for perinatal asphyxia on childhood outcomes. N Engl J Med. 2014;371:(2):140–9.

11. Wassink G, Davidson JO, Lear CA, Juul SE, Northington F, Bennet L, Gunn AJ. A working model for hypothermic neuroprotection. J Physiol. 2018;596:(23):5641–54.

12. Millar LJ, Shi L, Hoerder-Suabedissen A, Molnar Z. Neonatal Hypoxia Ischaemia: Mechanisms, Models, and Therapeutic Challenges. Front Cell Neurosci. 2017;11:78.

13. Vutskits L. Cerebral blood flow in the neonate. Paediatr Anaesth. 2014;24:(1):22–9.

14. Giussani DA. The fetal brain sparing response to hypoxia: physiological mechanisms. J Physiol. 2016;594:(5):1215–30.

15. Lear CA, Wassink G, Westgate JA, Nijhuis JG, Ugwumadu A, Galinsky R, Bennet L, Gunn AJ. The peripheral chemoreflex: indefatigable guardian of fetal physiological adaptation to labour. J Physiol. 2018;596:(23):5611–23.

16. Hassell KJ, Ezzati M, Alonso-Alconada D, Hausenloy DJ, Robertson NJ. New horizons for newborn brain protection: enhancing endogenous neuroprotection. Arch Dis Child Fetal Neonatal Ed. 2015;100:(6):F541–F552.

17. Dhillon SK, Lear CA, Galinsky R, Wassink G, Davidson JO, Juul S, Robertson NJ, Gunn AJ, Bennet L. The fetus at the tipping point: modifying the outcome of fetal asphyxia. J Physiol. 2018;596:(23):5571–92.

18. Mallard C, Vexler ZS. Modeling Ischemia in the Immature Brain: How Translational Are Animal Models? Stroke. 2015;46:(10):3006–11.

19. Semple BD, Blomgren K, Gimlin K, Ferriero DM, Noble-Haeusslein LJ. Brain development in rodents and humans: Identifying benchmarks of maturation and vulnerability to injury across species. Prog Neurobiol. 2013;

20. Sedmak G, Jovanov-Milosevic N, Puskarjov M, Ulamec M, Kruslin B, Kaila K, Judas M. Developmental Expression Patterns of KCC2 and Functionally Associated Molecules in the Human Brain. Cereb Cortex. 2016;26:(12):4574–89.

21. Workman AD, Charvet CJ, Clancy B, Darlington RB, Finlay BL. Modeling transformations of neurodevelopmental sequences across mammalian species. J Neurosci. 2013;33:(17):7368–83.

22. Rumajogee P, Bregman T, Miller SP, Yager JY, Fehlings MG. Rodent Hypoxia-Ischemia Models for Cerebral Palsy Research: A Systematic Review. Front Neurol. 2016;7:57.

23. Lupien SJ, McEwen BS, Gunnar MR, Heim C. Effects of stress throughout the lifespan on the brain, behaviour and cognition. Nat Rev Neurosci. 2009;10:(6):434–45.

24. Spoljaric A, Seja P, Spoljaric I, Virtanen MA, Lindfors J, Uvarov P, Summanen M, Crow AK, Hsueh B, Puskarjov M, Ruusuvuori E, Voipio J, Deisseroth K, Kaila K. Vasopressin excites interneurons to suppress hippocampal network activity across a broad span of brain maturity at birth. Proc Natl Acad Sci U S A. 2017;114:(50):E10819–E10828.

25. Morrison JL, Botting KJ, Darby JRT, David AL, Dyson RM, Gatford KL, Gray C, Herrera EA, Hirst JJ, Kim B, Kind KL, Krause BJ, Matthews SG, Palliser HK, Regnault TRH, Richardson BS, Sasaki A, Thompson LP, Berry MJ. Guinea pig models for translation of the developmental origins of health and disease hypothesis into the clinic. J Physiol. 2018;596:(23):5535–69.

26. Lear CA, Galinsky R, Wassink G, Yamaguchi K, Davidson JO, Westgate JA, Bennet L, Gunn AJ. The myths and physiology surrounding intrapartum decelerations: the critical role of the peripheral chemoreflex. J Physiol. 2016;594:(17):4711–25.

27. Schuchmann S, Schmitz D, Rivera C, Vanhatalo S, Salmen B, Mackie K, Sipilä ST, Voipio J, Kaila K. Experimental febrile seizures are precipitated by a hyperthermia-induced respiratory alkalosis. Nat Med. 2006;12:(7):817–23.

28. Ruusuvuori E, Kirilkin I, Pandya N, Kaila K. Spontaneous network events driven by depolarizing GABA action in neonatal hippocampal slices are not attributable to deficient mitochondrial energy metabolism. J Neurosci. 2010;30:(46):15638–42.

29. Ruusuvuori E, Kaila K: Carbonic anhydrases and brain pH in the control of neuronal excitability, Carbonic Anhydrase: Mechanism, Regulation, Links to Disease, and Industrial Applications. Edited by Frost S, McKenna R. Springer, 2014, pp 271–90

30. Pasternack M, Smirnov S, Kaila K. Proton modulation of functionally distinct GABAA receptors in acutely isolated pyramidal neurons of rat hippocampus. Neuropharmacology. 1996;35:(9-10):1279–88.

31. Wilkins ME, Hosie AM, Smart TG. Proton modulation of recombinant GABA(A) receptors: influence of GABA concentration and the beta subunit TM2-TM3 domain. J Physiol. 2005;567:(Pt 2):365–77.

32. Jensen FE, Applegate CD, Holtzman D, Belin TR, Burchfiel JL. Epileptogenic effect of hypoxia in the immature rodent brain. Ann Neurol. 1991;29:(6):629–37.

33. Peng BW, Justice JA, He XH, Sanchez RM. Decreased A-currents in hippocampal dentate granule cells after seizure-inducing hypoxia in the immature rat. Epilepsia. 2013;54:(7):1223–31.

34. Sampath D, White AM, Raol YH. Characterization of neonatal seizures in an animal model of hypoxic-ischemic encephalopathy. Epilepsia. 2014;55:(7):985–93.

35. Zanelli S, Goodkin HP, Kowalski S, Kapur J. Impact of transient acute hypoxia on the developing mouse EEG. Neurobiol Dis. 2014;68:37–46.

36. Zanelli SA, Rajasekaran K, Grosenbaugh DK, Kapur J. Increased excitability and excitatory synaptic transmission during in vitro ischemia in the neonatal mouse hippocampus. Neuroscience. 2015;310:279–89.

37. Saugstad OD. Resuscitation of newborn infants: from oxygen to room air. Lancet. 2010;376:(9757):1970–1.

38. Sun H, Juul HM, Jensen FE. Models of hypoxia and ischemia-induced seizures. J Neurosci Methods. 2016;260:252–60.

39. Helmy MM, Tolner EA, Vanhatalo S, Voipio J, Kaila K. Brain alkalosis causes birth asphyxia seizures, suggesting therapeutic strategy. Ann Neurol. 2011;69:(3):493–500.

40. Sanders RD, Manning HJ, Robertson NJ, Ma D, Edwards AD, Hagberg H, Maze M. Preconditioning and postinsult therapies for perinatal hypoxic-ischemic injury at term. Anesthesiology. 2010;113:(1):233–49.

41. Pappas A, Shankaran S, Laptook AR, Langer JC, Bara R, Ehrenkranz RA, Goldberg RN, Das A, Higgins RD, Tyson JE, Walsh MC. Hypocarbia and adverse outcome in neonatal hypoxic-ischemic encephalopathy. J Pediatr. 2011;158:(5):752–8.

42. Reynolds RD, Pilcher J, Ring A, Johnson R, McKinley P. The Golden Hour: care of the LBW infant during the first hour of life one unit’s experience. Neonatal Netw. 2009;28:(4):211–9.

43. Shah V, Hodgson K, Seshia M, Dunn M, Schmolzer GM. Golden hour management practices for infants <32 weeks gestational age in Canada. Paediatr Child Health. 2018;23:(4):e70–e76.

44. McNamara HM, Dildy GA. Continuous intrapartum pH, pO2, pCO2, and SpO2 monitoring. Obstet Gynecol Clin North Am. 1999;26:(4):671–93.

45. Voipio J: Diffusion and buffering aspects of H+, HCO3-, and CO2 movements in brain tissue, pH and brain function. Edited by Kaila K, Ransom BR. Wiley-Liss, 1998, pp 45–66

46. Chesler M. Regulation and modulation of pH in the brain. Physiol Rev. 2003;83:(4):1183–221.

47. Summanen M, Back S, Voipio J, Kaila K. Surge of Peripheral Arginine Vasopressin in a Rat Model of Birth Asphyxia. Front Cell Neurosci. 2018;12:2.

48. Woody CD, Marshall WH, Besson JM, Thompson HK, Aleonard P, Albe-Fessard D. Brain potential shift with respiratory acidosis in the cat and monkey. Am J Physiol. 1970;218:(1):275–83.

49. Voipio J, Tallgren P, Heinonen E, Vanhatalo S, Kaila K. Millivolt-scale DC shifts in the human scalp EEG: Evidence for a nonneuronal generator. J Neurophysiol. 2003;89:(4):2208–14.

50. Collins JA, Rudenski A, Gibson J, Howard L, O’Driscoll R. Relating oxygen partial pressure, saturation and content: the haemoglobin-oxygen dissociation curve. Breathe (Sheff). 2015;11:(3):194–201.

51. Clancy B, Finlay BL, Darlington RB, Anand KJ. Extrapolating brain development from experimental species to humans. Neurotoxicology. 2007;28:(5):931–7.

52. Erecinska M, Cherian S, Silver IA. Energy metabolism in mammalian brain during development. Prog Neurobiol. 2004;73:(6):397–445.

53. Plate KH. Mechanisms of angiogenesis in the brain. J Neuropathol Exp Neurol. 1999;58:(4):313–20.

54. Hirst JJ, Palliser HK, Shaw JC, Crombie G, Walker DW, Zakar T. Birth and Neonatal Transition in the Guinea Pig: Experimental Approaches to Prevent Preterm Birth and Protect the Premature Fetus. Front Physiol. 2018;9:1802.

55. Turek Z, Ringnalda BE, Moran O, Kreuzer F. Oxygen transport in guinea pigs native to high altitude (Junin, Peru, 4,105 m). Pflugers Arch. 1980;384:(2):109–15.

56. Gonzalez-Obeso E, Docio I, Olea E, Cogolludo A, Obeso A, Rocher A, Gomez-Nino A. Guinea Pig Oxygen-Sensing and Carotid Body Functional Properties. Front Physiol. 2017;8:285.

57. Lam TI, Wise PM, O’Donnell ME. Cerebral microvascular endothelial cell Na/H exchange: evidence for the presence of NHE1 and NHE2 isoforms and regulation by arginine vasopressin. Am J Physiol Cell Physiol. 2009;297:(2):C278–C289.

58. Helmy MM, Ruusuvuori E, Watkins PV, Voipio J, Kanold PO, Kaila K. Acid extrusion via blood-brain barrier causes brain alkalosis and seizures after neonatal asphyxia. Brain. 2012;135:(Pt 11):3311–9.

59. Mitsufuji N, Yoshioka H, Tominaga M, Okano S, Nishiki T, Sawada T. Intracellular alkalosis during hypoxia in newborn mouse brain in the presence of systemic acidosis: a phosphorus magnetic resonance spectroscopic study. Brain Dev. 1995;17:(4):256–60.

60. Ala-Kurikka, T., Summanen, M., Pospelov, A. S., Alafuzoff, A., Voipio, J., and Kaila, K. A novel rat model of term intrapartum asphyxia. 3^rd^ Nordic Neuroscience Meeting. Session A 02. 2019. 12-6-2019.

61. Lagercrantz H, Bistoletti P. Catecholamine release in the newborn infant at birth. Pediatr Res. 1977;11:(8):889–93.

62. Evers KS, Wellmann S. Arginine Vasopressin and Copeptin in Perinatology. Front Pediatr. 2016;4:75.

63. Hankins GD, Speer M. Defining the pathogenesis and pathophysiology of neonatal encephalopathy and cerebral palsy. Obstet Gynecol. 2003;102:(3):628–36.

64. Malin GL, Morris RK, Khan KS. Strength of association between umbilical cord pH and perinatal and long term outcomes: systematic review and meta-analysis. BMJ. 2010;340:c1471.

65. Belai Y, Goodwin TM, Durand M, Greenspoon JS, Paul RH, Walther FJ. Umbilical arteriovenous PO2 and PCO2 differences and neonatal morbidity in term infants with severe acidosis. Am J Obstet Gynecol. 1998;178:(1 Pt 1):13–9.

66. Williams KP, Singh A. The correlation of seizures in newborn infants with significant acidosis at birth with umbilical artery cord gas values. Obstet Gynecol. 2002;100:(3):557–60.

67. Dyson RM, Palliser HK, Kelleher MA, Hirst JJ, Wright IM. The guinea pig as an animal model for studying perinatal changes in microvascular function. Pediatr Res. 2012;71:(1):20–4.

68. Pearson T, Nuritova F, Caldwell D, Dale N, Frenguelli BG. A depletable pool of adenosine in area CA1 of the rat hippocampus. J Neurosci. 2001;21:(7):2298–307.

69. Lutas A, Birnbaumer L, Yellen G. Metabolism regulates the spontaneous firing of substantia nigra pars reticulata neurons via KATP and nonselective cation channels. J Neurosci. 2014;34:(49):16336–47.

70. Lynch NE, Stevenson NJ, Livingstone V, Murphy BP, Rennie JM, Boylan GB. The temporal evolution of electrographic seizure burden in neonatal hypoxic ischemic encephalopathy. Epilepsia. 2012;53:(3):549–57.

71. Lee J, Taira T, Pihlaja P, Ransom BR, Kaila K. Effects of CO2 on excitatory transmission apparently caused by changes in intracellular pH in the rat hippocampal slice. Brain Res. 1996;706:(2):210–6.

72. Schuchmann S, Hauck S, Henning S, Gruters-Kieslich A, Vanhatalo S, Schmitz D, Kaila K. Respiratory alkalosis in children with febrile seizures. Epilepsia. 2011;52:(11):1949–55.

73. Tolner EA, Hochman DW, Hassinen P, Otahal J, Gaily E, Haglund MM, Kubova H, Schuchmann S, Vanhatalo S, Kaila K. Five percent CO(2) is a potent, fast-acting inhalation anticonvulsant. Epilepsia. 2011;52:(1):104–14.

74. Seidl R, Stockler-Ipsiroglu S, Rolinski B, Kohlhauser C, Herkner KR, Lubec B, Lubec G. Energy metabolism in graded perinatal asphyxia of the rat. Life Sci. 2000;67:(4):421–35.

75. Agostoni A, Stabilini R, Bernasconi C, Gerli GC. The oxyhemoglobin dissociation curve in hypercapnic patients. Am Heart J. 1974;87:(5):670–2.

76. Jensen FB. Red blood cell pH, the Bohr effect, and other oxygenation-linked phenomena in blood O2 and CO2 transport. Acta Physiol Scand. 2004;182:(3):215–27.

77. Yli BM, Kjellmer I. Pathophysiology of foetal oxygenation and cell damage during labour. Best Pract Res Clin Obstet Gynaecol. 2016;30:9–21.

78. Bennet L, Westgate JA, Liu YC, Wassink G, Gunn AJ. Fetal acidosis and hypotension during repeated umbilical cord occlusions are associated with enhanced chemoreflex responses in near-term fetal sheep. J Appl Physiol (1985). 2005;99:(4):1477–82.

79. Akca O, Sessler DI, Delong D, Keijner R, Ganzel B, Doufas AG. Tissue oxygenation response to mild hypercapnia during cardiopulmonary bypass with constant pump output. Br J Anaesth. 2006;96:(6):708–14.

80. Ratnaraj J, Kabon B, Talcott MR, Sessler DI, Kurz A. Supplemental oxygen and carbon dioxide each increase subcutaneous and intestinal intramural oxygenation. Anesth Analg. 2004;99:(1):207–11.

81. Davis JW. The relationship of base deficit to lactate in porcine hemorrhagic shock and resuscitation. J Trauma. 1994;36:(2):168–72.

82. Chawla LS, Nader A, Nelson T, Govindji T, Wilson R, Szlyk S, Nguyen A, Junker C, Seneff MG. Utilization of base deficit and reliability of base deficit as a surrogate for serum lactate in the peri-operative setting. BMC Anesthesiol. 2010;10:16.

83. Hood VL, Tannen RL. Protection of acid-base balance by pH regulation of acid production. N Engl J Med. 1998;339:(12):819–26.

84. Szakmar E, Jermendy A, El-Dib M. Respiratory management during therapeutic hypothermia for hypoxic-ischemic encephalopathy. J Perinatol. 2019;39:(6):763–73.

85. Vannucci RC, Towfighi J, Heitjan DF, Brucklacher RM. Carbon dioxide protects the perinatal brain from hypoxic-ischemic damage: an experimental study in the immature rat. Pediatrics. 1995;95:(6):868–74.

86. Ikonen RS, Janas MO, Koivikko MJ, Laippala P, Kuusinen EJ. Hyperbilirubinemia, hypocarbia and periventricular leukomalacia in preterm infants: relationship to cerebral palsy. Acta Paediatr. 1992;81:(10):802–7.

87. Wiswell TE, Graziani LJ, Kornhauser MS, Stanley C, Merton DA, McKee L, Spitzer AR. Effects of hypocarbia on the development of cystic periventricular leukomalacia in premature infants treated with high-frequency jet ventilation. Pediatrics. 1996;98:(5):918–24.

88. Thome UH, Ambalavanan N. Permissive hypercapnia to decrease lung injury in ventilated preterm neonates. Semin Fetal Neonatal Med. 2009;14:(1):21–7.

89. Ryu J, Haddad G, Carlo WA. Clinical effectiveness and safety of permissive hypercapnia. Clin Perinatol. 2012;39:(3):603–12.

90. Logan JW. First, Do No Harm. Consequences of Permissive Hypercapnia in the Neonate. Respir Care. 2018;63:(8):1070–2.

91. Kaiser JR, Gauss CH, Williams DK. The effects of hypercapnia on cerebral autoregulation in ventilated very low birth weight infants. Pediatr Res. 2005;58:(5):931–5.

92. Klinger G, Beyene J, Shah P, Perlman M. Do hyperoxaemia and hypocapnia add to the risk of brain injury after intrapartum asphyxia? Arch Dis Child Fetal Neonatal Ed. 2005;90:(1):F49–F52.

93. Hillered L, Ernster L, Siesjo BK. Influence of in vitro lactic acidosis and hypercapnia on respiratory activity of isolated rat brain mitochondria. J Cereb Blood Flow Metab. 1984;4:(3):430–7.

94. Li J, Hoskote A, Hickey C, Stephens D, Bohn D, Holtby H, Van AG, Redington AN, Adatia I. Effect of carbon dioxide on systemic oxygenation, oxygen consumption, and blood lactate levels after bidirectional superior cavopulmonary anastomosis. Crit Care Med. 2005;33:(5):984–9.

95. Jensen EC, Bennet L, Hunter CJ, Power GC, Gunn AJ. Post-hypoxic hypoperfusion is associated with suppression of cerebral metabolism and increased tissue oxygenation in near-term fetal sheep. J Physiol. 2006;572:(Pt 1):131–9.

96. Saugstad OD, Oei JL, Lakshminrusimha S, Vento M. Oxygen therapy of the newborn from molecular understanding to clinical practice. Pediatr Res. 2019;85:(1):20–9.

97. Albertine KH. Utility of large-animal models of BPD: chronically ventilated preterm lambs. Am J Physiol Lung Cell Mol Physiol. 2015;308:(10):L983–L1001.

98. Laffey JG, Tanaka M, Engelberts D, Luo X, Yuan S, Tanswell AK, Post M, Lindsay T, Kavanagh BP. Therapeutic hypercapnia reduces pulmonary and systemic injury following in vivo lung reperfusion. Am J Respir Crit Care Med. 2000;162:(6):2287–94.

99. Curley GF, Laffey JG, Kavanagh BP. CrossTalk proposal: there is added benefit to providing permissive hypercapnia in the treatment of ARDS. J Physiol. 2013;591:(11):2763–5.

100. Currin RT, Gores GJ, Thurman RG, Lemasters JJ. Protection by acidotic pH against anoxic cell killing in perfused rat liver: evidence for a pH paradox. FASEB J. 1991;5:(2):207–10.

101. Halestrap AP. Calcium-dependent opening of a non-specific pore in the mitochondrial inner membrane is inhibited at pH values below 7. Implications for the protective effect of low pH against chemical and hypoxic cell damage. Biochem J. 1991;278 (Pt 3):715–9.

102. Andrienko TN, Pasdois P, Pereira GC, Ovens MJ, Halestrap AP. The role of succinate and ROS in reperfusion injury - A critical appraisal. J Mol Cell Cardiol. 2017;110:1–14.

103. Nita DA, Vanhatalo S, Lafortune FD, Voipio J, Kaila K, Amzica F. Nonneuronal origin of CO_2_-related DC EEG shifts: an in vivo study in the cat. J Neurophysiol. 2004;92:(2):1011–22.

104. Kiviniemi V, Korhonen V, Kortelainen J, Rytky S, Keinanen T, Tuovinen T, Isokangas M, Sonkajarvi E, Siniluoto T, Nikkinen J, Alahuhta S, Tervonen O, Turpeenniemi-Hujanen T, Myllyla T, Kuittinen O, Voipio J. Real-time monitoring of human blood-brain barrier disruption. PLoS One. 2017;12:(3):e0174072.

